# Loneliness and the social brain: how perceived social isolation impairs human interactions

**DOI:** 10.1101/2021.03.03.433569

**Authors:** Jana Lieberz, Simone G. Shamay-Tsoory, Nira Saporta, Timo Esser, Ekaterina Kuskova, Birgit Stoffel-Wagner, René Hurlemann, Dirk Scheele

## Abstract

Loneliness is a painful condition associated with increased risk for premature mortality. The formation of new, positive social relationships can alleviate feelings of loneliness, but requires rapid trustworthiness decisions during initial encounters and it is still unclear how loneliness hinders interpersonal trust. Here, we use a multimodal approach including behavioral, psychophysiological, hormonal, and neuroimaging measurements to probe a trust-based mechanism underlying impaired social interactions in loneliness. Pre-stratified healthy individuals with high loneliness scores (*n* = 42 out of a screened sample of 3678 adults) show reduced oxytocinergic and affective responsiveness to a positive conversation, report less interpersonal trust, and prefer larger social distances compared to controls (*n* = 40). Moreover, lonely individuals are rated as less trustworthy compared to controls and identified by the blinded confederate better than chance. During initial trust decisions, lonely individuals exhibit attenuated limbic and striatal activation and blunted functional connectivity between the anterior insula and occipitoparietal regions, which correlates with the diminished affective responsiveness to the positive social interaction. This neural response pattern is not mediated by loneliness-associated psychological symptoms. Thus, our results indicate compromised integration of trust-related information as a shared neurobiological component in loneliness, yielding a reciprocally reinforced trust bias in social dyads.

## 1. Introduction

Humans are an essentially social species with the motivation to form and maintain interpersonal relationships as a fundamental organizational principle of behavior. When a person’s need to belong is not satisfied, distressful feelings of loneliness, that is perceived social isolation, occur. Various lines of research indicate that loneliness has detrimental effects on mental and physical health, evident in increased risk of psychological disorders, cognitive decline, and all-cause mortality.^[1,2]^ As such, loneliness has been identified as a public health challenge with prevalence rates up to 33 % across age,^[3]^ but the unclear etiological mechanisms leading to and fostering the maintenance of loneliness hamper the development of neurobiologically-informed interventions not only on the individual but also the societal level.^[4-6]^

From an evolutionary perspective, loneliness may have evolved to motivate the formation of new social relationships, in the same way as hunger induces scavenging.^[7-9]^ However, when the connection with other individuals fails, loneliness impairs inflammatory and immune responses^[6,10]^ and promotes a phenotypic hypersensitivity to social threats and self-centered behavior.^[8,11]^ The perception of the social environment as threatening may lead to various negative biases in loneliness. For instance, it has been suggested that lonely individuals allocate their attention faster towards threatening social stimuli, anticipate rejection more often, and exhibit negative attribution styles.^[11]^ Eventually, even positive social interactions might fail to alleviate feelings of loneliness, as lonely individuals show reduced positive ratings of social encounters and attenuated reward-associated brain activity in response to positive social stimuli.^[12,13]^ Importantly, however, while the detrimental impact of loneliness on social interactions is well established and theoretical frameworks point to negative biases and selfish behavior as putative mediators, the neurobiological mechanisms that hinder the formation of new, positive relationships and thus the alleviation of loneliness are still elusive.

In human societies, the development of positive relationships is based mainly on cooperation, with non-cooperative behaviors evoking avoidance or even punishment. However, during initial encounters, when there is no prior information about the likelihood of reciprocity, rapid trustworthiness decisions are required for the formation of new relationships. Importantly, preliminary evidence indicates that interpersonal trust is reduced in lonely individuals.^[11]^ In addition, the neural circuits of trust and loneliness are largely intertwined and share neuroanatomical pathways via the amygdala, the anterior insula (AI), the medial prefrontal cortex (mPFC), the nucleus accumbens (NAcc), and the temporoparietal junction (TPJ).^[6,14-17]^ Nevertheless, it is still unclear whether these brain regions might contribute to reduced interpersonal trust in loneliness, as they have been associated with various cognitive processes.^[18,19]^ This would indicate that the selectivity of activation is low and specific inferences are not valid without evidence that the assumed process (i.e. interpersonal trust) is engaged.^[20]^

Thus, the current study aims to examine to what extent loneliness relates to interpersonal trust, and whether activity and connectivity of the aforementioned neural circuit would be altered in lonely individuals during situations that specifically require trustworthiness decisions. We hypothesized that participants with high loneliness scores (high-lonely, HL) would exhibit diminished interpersonal trust in self-report and behavioral measurements as well as altered trust-associated brain activity and connectivity. Furthermore, given the key role of interpersonal trust for the development of positive relationships, we hypothesized that reduced interpersonal trust and its underlying brain activity would mechanistically contribute to attenuated benefits from a positive social interaction in lonely individuals.

To test our hypotheses, we implemented a multimodal pre-stratification approach including behavioral, psychophysiological, hormonal, and neuroimaging measurements. We screened a sample of *n* = 3678 individuals and included *n* = 42 HL and *n* = 40 controls (low-lonely, LL) who participated in a positive conversation with an unfamiliar confederate and underwent functional magnetic resonance imaging (fMRI) during which they played an adapted version of the well-established trust game.^[21]^ Specifically, we hypothesized increased positive and decreased negative mood ratings as response to the positive conversation across all participants. Moreover, we expected that affective responses would be reduced in HL participants. In contrast, we hypothesized that HL and LL participants would not differ regarding their physiological responsiveness to the conversation (i.e. changes in electrodermal activity and heart rate), as we assumed that the reduced affective responsiveness would be based on negative biases rather than differences in physiological arousal.

To probe the hypotheses of reduced interpersonal trust and trust-associated brain activity as a potential mechanism underlying the impaired reactivity to social interactions in loneliness, we first measured self-reported interpersonal trust and the ideal and uncomfortable interpersonal distance during a stop-approach paradigm (cf.^[22]^) as behavioral measurement of interpersonal trust towards the confederate. We then contrasted brain activity during the fMRI trust game with a risk game control condition to test the hypothesized altered brain activity in the amgydala, AI, mPFC, NAcc, and TPJ and to further explore whether differences in brain activity would be accompanied by altered functional connectivity. We lastly hypothesized that the observed differences in responsiveness to the positive social interaction of HL compared to LL participants would correlate with the trust assessments. We controlled for the influence of possible confounding variables such as depressive symptomatology, social anxiety, and childhood maltreatment.

In addition to these hypotheses, we assessed further exploratory variables to better characterize the response profile to the positive social interaction in lonely individuals. We collected saliva samples before and after the task to explore hormonal and immunological reactivity. Salivary assessments consisted of the hypothalamic peptide oxytocin, which is crucially involved in human bonding and trust,^[23-25]^ as well as cortisol and immunoglobulin A (IgA) concentrations as markers of stress and immune system responses^[26]^ to the social interaction, in addition to baseline immune parameters in blood. Moreover, the blinded confederate in the social interaction task estimated the group affiliation (HL versus LL) and rated the trustworthiness of the participants to examine the social transmission of loneliness. Finally, as sex differences in the neural correlates of loneliness have been identified recently,^[14]^ we conducted moderator analyses to explore the potential influence of the participants’ sex on loneliness effects in our sample.

## 2. Results

### 2.1. Loneliness and impaired social interaction

First, we examined behavioral, hormonal, and psychophysiological responses to a positive, real-life social interaction in a controlled setting. As expected, across groups (HL: *n* = 42, 21 female; LL: *n* = 40, 20 female, cf. **Table S1**), the positive interaction was experienced as very pleasant [*M* ± *SD*: 82.19 ± 16.73 on a visual analogue scale (VAS) ranging from 0 (“not pleasant at all”) to 100 (“very pleasant”); see **Figure 1A**] and significantly increased positive mood: specifically, we observed an increase in positive affect and in vigor [for all main effects of time (before versus after the interaction): *F*s > 11.06, *P*s < 0.002, η_p_ ^2^ > 0.12; 95 % confidence interval (CI) of increase in scores of the positive affect: 1.25 to 3.30; vigor: 0.96 to 3.85]. An increase in general physiological activity was evident for the skin conductance level (SCL) and heart beats per minute (BPM) (main effect of time for SCL: *F*(_1,72)_ = 5.89, *P* = 0.018, η_p_^2^ = 0.08, 95 % CI of increase: 2.10 to 2.88 μS, see **Figure 1B;** BPM: *F*_(1,70)_ = 11.36, *P* = 0.001, η_p_^2^ = 0.14, 95 % CI: 3.56 to 5.47 BPM, see **Figure 1C**). Furthermore, the positive social interaction led to elevated salivary oxytocin and IgA levels [area under the curve (AUC_I_) describing the increase tested against zero: all *t*s > 2.59, *P*s < 0.013, *d*s > 0.38; 95 % CI of AUC_I_ of oxytocin: 1.63 to 5.62; IgA: 0.06 to 0.51; mean percentage increase between before and after the interaction ± SD in oxytocin: 17.03 ± 31.23 %; IgA: 16.21 ± 47.52 %].

**Figure 1.**
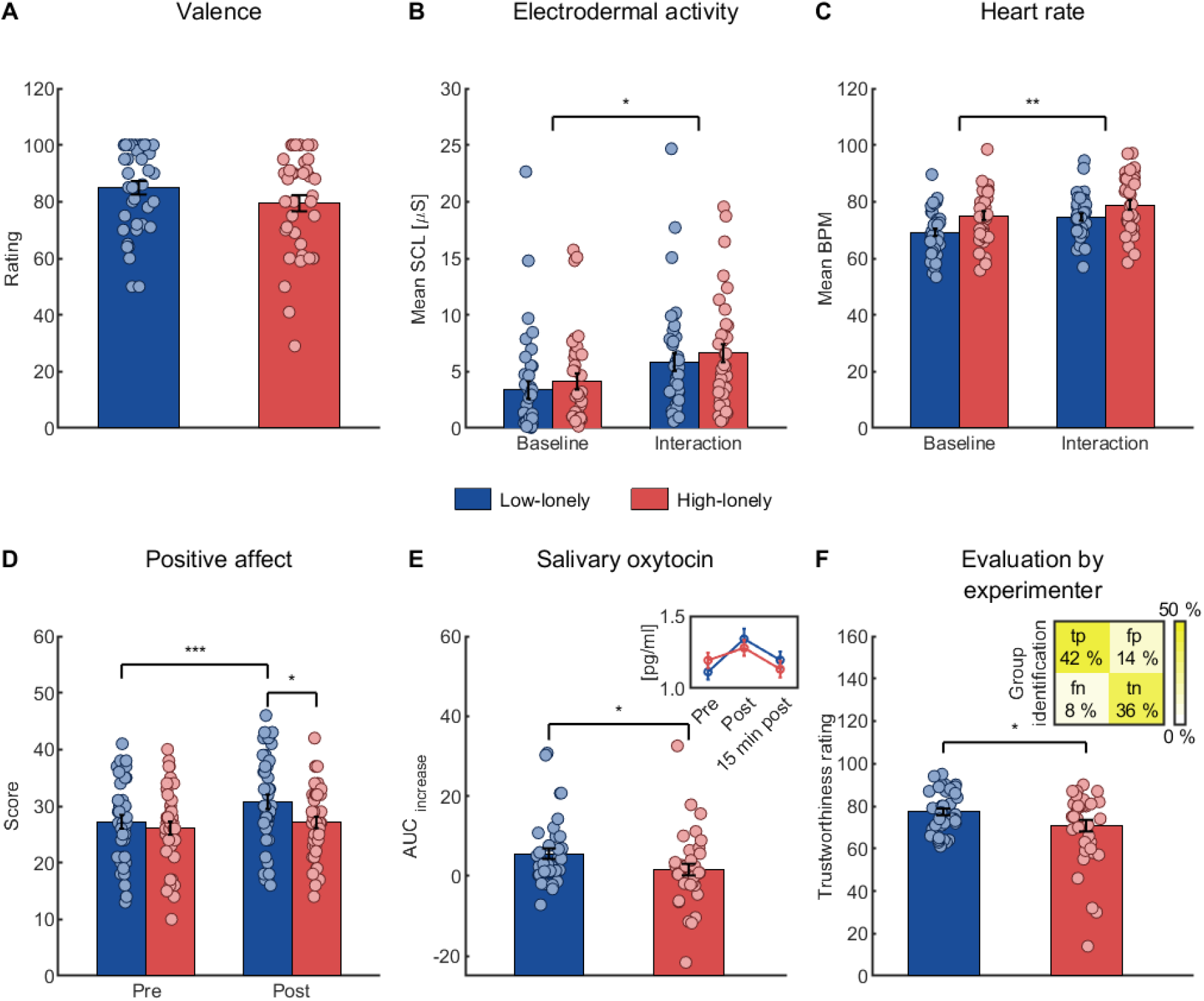
Response profile to the positive social interaction paradigm. Participants rated the positive social interaction as very pleasant on a visual analogue scale (VAS) ranging from 0 (“not pleasant at all”) to 100 (“very pleasant”) and ratings did not differ between groups (**A**). Across groups, mean skin conductance level (SCL; **B**) and mean heart beats per minute (BPM) increased during the social interaction compared to a 5-minute rest baseline (**C**). However, high-lonely participants showed diminished reactivity to the social interaction. Positive affect increased in low-lonely but not high-lonely participants (**D**) and the area under the curve (AUC) measuring the increase in salivary oxytocin levels was attenuated in the high-lonely sample (**E**). The inlay displays the group mean salivary oxytocin concentration for each time point. After completion of the social interaction, the experimenter rated high-lonely participants as less trustworthy on a VAS ranging from 0 (“not trustworthy at all”) to 100 (“very trustworthy”) and identified high-lonely participants significantly better than by chance (**F**). The inlay displays the percentage of false negative (fn; high-lonely classified as low-lonely), false positive (fp; low-lonely classified as high-lonely), true negative (tn; low-lonely classified as low-lonely), and true positive (tp; high-lonely classified as high-lonely) classifications. All bars represent group means. Error bars indicate standard errors of the mean. Dots are jittered for purposes of presentation. * P < 0.05, ** P < 0.01, *** P < 0.001.

Importantly, as hypothesized, HL participants exhibited attenuated self-reported affective reactivity to the positive interaction (interaction of time and group for positive affect: *F*_(1,77)_ = 6.43, *P* = 0.013, η_p2_ = 0.08). Post-hoc *t*-tests revealed a significant increase in positive affect in LL participants [*t*_(39)_ = 5.02, *P* < 0.0001 after Bonferroni-correction (*P*_cor_), *d* = 0.45, 95 % CI of increase in score: 2.10 to 4.95], but not in HL participants (*t*_(38)_ = 1.42, *P*_cor_ = 0.658, 95 % CI: -0.43 to 2.43, see **Figure 1D**). By contrast, the physiological reactivity to the positive social interaction did not differ between groups (no significant interaction of time with group for SCL or BPM measurements, all *P*s > 0.075), suggesting that observed affective group effects were not based on differences in the experiences of physiological arousal.

Interestingly, we did not observe baseline differences in plasma (*t*_(77)_ = 0.13, *P* = 0.895, 95 % CI of group difference: -0.42 to 0.48 pg/ml) or salivary oxytocin levels (*t*_(76)_ = 1.09, *P* = 0.278, 95 % CI: -0.07 to 0.23 pg/ml), but HL participants showed a reduced increase in salivary oxytocin levels compared to LL participants (*t*_(75)_ = -2.04, *P* = 0.045, *d* = -0.47, 95 % CI of AUC_I_ difference between groups: -7.93 to -0.10; see **Figure 1E**). Consistent with the notion that loneliness can be perceived by others ^[27]^, the blinded experimenters were significantly better than chance in identifying HL participants after the interaction (78 % correct, χ^2^_(1)_ = 24.82, *P* < 0.0001; specificity: 72 %; sensitivity: 85 %). In addition, the experimenters rated HL participants as less trustworthy than LL individuals (*t*_(61.13)_ = -2.06, *P* = 0.043, *d* = -0.47, 95 % CI of group difference: -12.82 to -0.20; see **Figure 1F**).

Collectively, we confirmed that HL participants showed not only a reduced responsiveness to the positive social interaction as evident for self-reported positive affect but also exhibited an attenuated oxytocinergic response. Furthermore, loneliness affected the experimenter’s perception of the participants. In the following, we examined the potential impact of interpersonal trust on the impaired social interaction effects in HL participants. For further analyses of the social interaction and immunology, see Supplementary Analyses, **Figure S1**, and **Table S2**.

### 2.2. Loneliness and reduced interpersonal trust

In line with our hypotheses, HL participants reported significantly less interpersonal trust compared to LL individuals (*t*_(80)_ = -4.62, *P* < 0.0001, *d* = -1.02, 95 % CI of group difference in scores: -0.79 to -0.31; see **Figure 2A**) and self-reported trust positively correlated with the positive affect after the positive social interaction (ρ_(78)_ = 0.28, *P* = 0.014, 95 % CI: 0.06 to 0.47). Reduced trust in loneliness was also evident in form of a greater preferred interpersonal distance to strangers. Mixed analyses of variance (ANOVA) with time (before versus after completing the positive social interaction paradigm) as within-subject factor and group (HL versus LL) as between subject factor yielded main effects of group for the ideal (*F*_(1,77)_ = 7.17, *P* = 0.009, η_p_^2^ = 0.09, 95 % CI of group difference: 0.03 to 0.20 m; see **Figure 2B**) and slightly uncomfortable distance (*F*_(1,77)_ = 4.05, *P* = 0.048, η_p_^2^ = 0.05, 95 % CI: 0.001 to 0.13 m). Although distances decreased after the positive interaction (main effect of time for the ideal distance: *F*_(1,77)_ = 41.63, *P* < 0.0001, η_p_^2^ = 0.35, 95 % CI of decrease: -0.13 to -0.07 m; uncomfortable distance: *F*_(1,77)_ = 5.94, *P* = 0.017, η_p_^2^ = 0.07, 95 % CI: -0.04 to -0.004 m), the positive interaction was not sufficient to alleviate group differences (all time with group interactions *P*s > 0.376). As expected, self-reported trust negatively correlated with the ideal distance (ρ_(78)_ = -0.24, *P* = 0.032, 95 % CI: -0.44 to -0.02; see **Figure 2C**) but not with the distance at which participants felt slightly uncomfortable (ρ_(78)_ = -0.08, *P* = 0.509, 95 % CI: -0.29 to 0.15).

**Figure 2.**
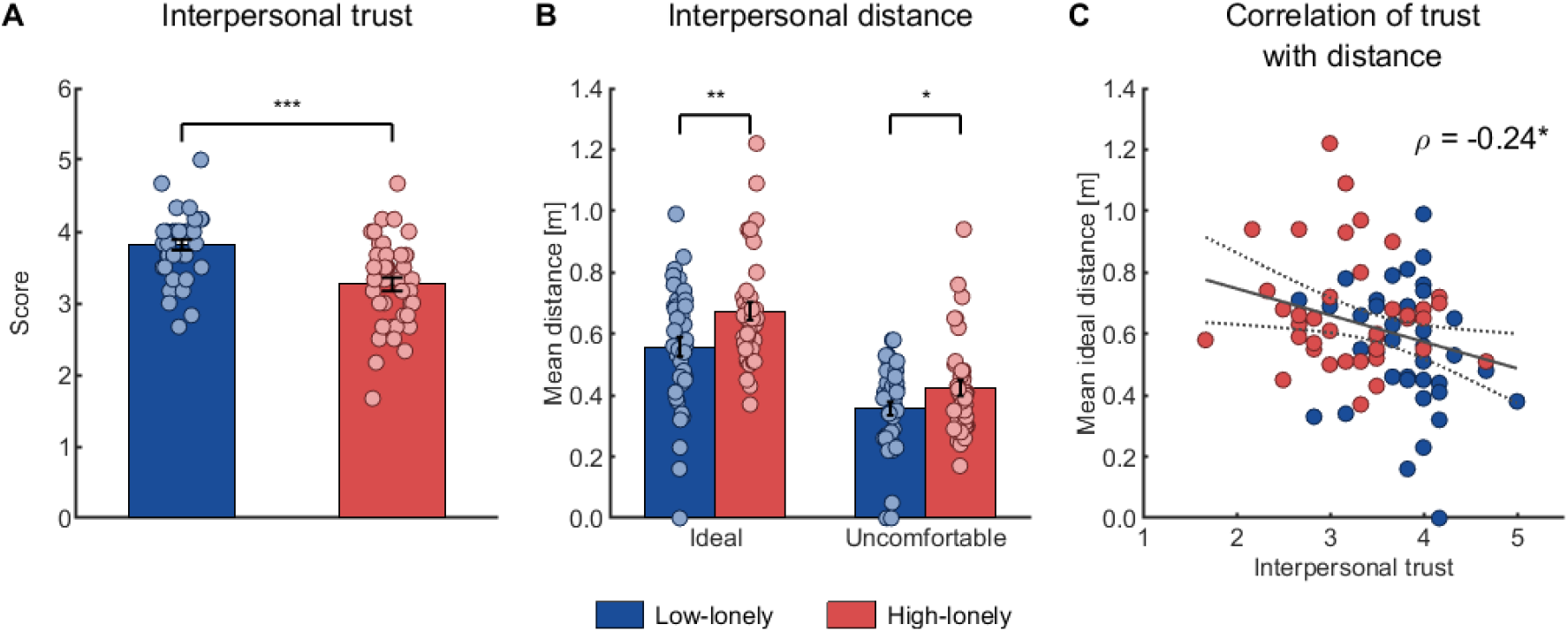
Reduced interpersonal trust and larger social distance in loneliness. High-lonely participants reported less interpersonal trust (A). Across time points, high-lonely participants stopped at a larger ideal and uncomfortable distance to the experimenter in the stop-distance paradigm (B). Across groups, self-reported interpersonal trust negatively correlated with the mean ideal distance of participants (C), that is individuals with lower interpersonal trust prefered a greater ideal interpersonal distance. The dashed line represents the 95%-confidence interval of the plotted regression line. All bars represent group means. Error bars indicate standard errors of the mean. Dots on bar plots are jittered for purposes of presentation. * P < 0.05, ** P < 0.01, *** P < 0.001.

We further analyzed investment behavior during the trust game by calculating a mixed ANOVA (within-subject factor: game type trust versus risk, between-subject factor: group). The HL subsample was characterized by overall lower investments (main effect of group: *F*_(1,63)_ = 4.01, *P* = 0.0495, η_p_^2^ = 0.06; 95 % CI of group difference: -2.24 to -0.002 €). Importantly, the absence of significant effects of game type (all *P*s > 0.119 for a main effect or interaction with group) indicates that the implemented risk game constitutes a well-matched control condition for the trust game, as our results show that potential neural differences between conditions cannot be related to different investment choices.

### 2.3. Loneliness and trust-related brain activity and connectivity

To investigate the association of loneliness with trust-related brain activity, we contrasted brain activity during the trust game to the risk game. In a first step, we confirmed that our implementation of the trust game led to enhanced trust-related brain activity. Whole-brain analyses indeed revealed significantly increased activity during the trust game compared to the risk game in several brain regions associated with trust including the insula, mPFC, hippocampus and amygdala, and TPJ [all *P*s < 0.05 on peak level after family-wise error (FWE) correction; see Supplementary Analyses and **Table S3** for details and further whole brain analyses]. We then examined group differences in the reactions to the trust game (trust game > risk game). HL participants showed significantly reduced trust-associated activity in the left AI (−26, 10, -18, *t*_(57)_ = 4.07, FWE-corrected *P* = 0.034; see **Figure 3A**), right NAcc (12, 8, -8, *t*_(57)_ = 2.88, FWE-corrected *P* = 0.031; see **Figure 3B**), and left amygdala (−20, -8, -16, *t*_(57)_ = 3.56, FWE-corrected *P* = 0.042; see **Figure 3C**). No significant opposite effects were observed (i.e. increased brain activity during the trust game in HL participants) and groups did not differ in trust-related mPFC or TPJ activity (all FWE-corrected *P*s ≥ 0.209). To further characterize the observed interaction of game type and group in the left amygdala, left AI, and right NAcc, we compared parameter estimates using two-sample *t*-tests for each cluster. Results revealed that HL participants did not differ from LL participants in game conditions *per se* (all *P*_cor_ > 0.072) but rather showed a blunted differentiation (i.e. smaller activity increase) between trust- and risk-related trials in brain regions associated with the evaluation of trustworthiness, risk of betrayal, and reward anticipation.^[15]^

**Figure 3.**
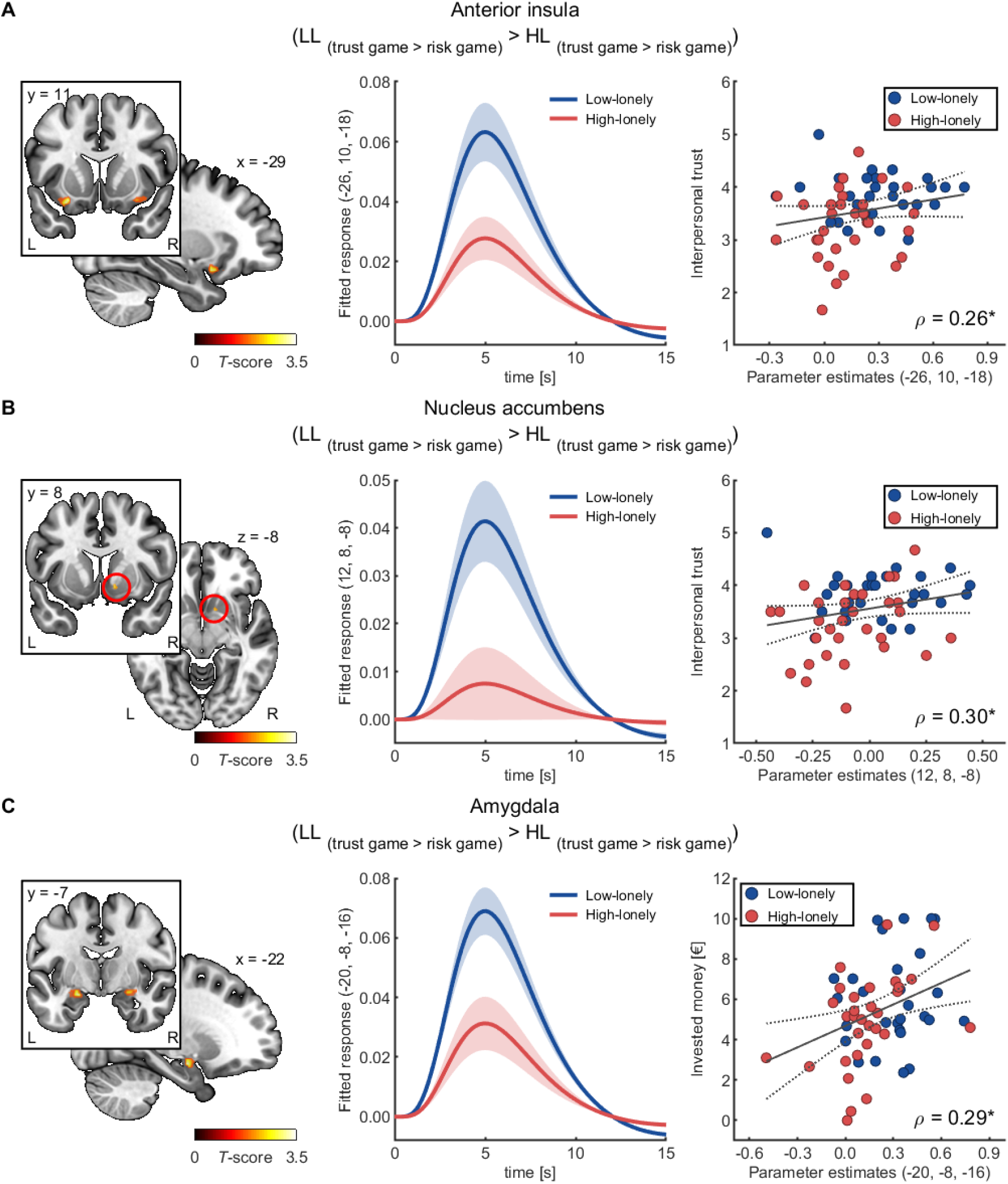
Reduced trust-associated brain activity in high-lonely participants. High-lonely participants exhibited less activity in the trust game relative to the risk game in the left anterior insula (**A**), the right nucleus accumbens (**B**), and the left amygdala (**C**). In the pooled sample, responses in the anterior insula and the right nucleus accumbens positively correlated with self-reported trust, while parameter estimates of the left amygdala activity during the trust game (compared to the risk game) were positively associated with the invested money across conditions. For illustration purpose clusters are shown with significance levels of P < 0.05 uncorrected. The shaded areas show the standard error of the mean of the estimated time courses based on the canonical hemodynamic response function as used in SPM12 multiplied by the parameter estimates of the trust game > risk game contrast. The dashed lines represent the 95%-confidence intervals of the plotted regression lines. Abbreviations: HL, high-lonely; L, left; LL, low-lonely; R, right. * P < 0.05.

To probe the robustness of the reduced trust-associated brain activity observed in HL participants, we further analyzed our data by conducting Bayesian inference analyses as implemented in SPM12. Results provide strong evidence that the AI activity is reduced in HL participants compared to LL participants [-26, 10, -18, log odds Bayes factor for attenuated activity in HL participants versus no group differences or enhanced activity in HL participants compared to controls (logBF) = 3.28]. Thus, the Bayes analyses confirm the results of the frequentist analyses for the left AI, but not for the amygdala or NAcc. Notably, our data also provide strong evidence for reduced mPFC activity that was not detected by the frequentist analyses (0, 52, 10, logBF = 3.62; for further results of the Bayesian analyses that exceed the predefined ROI, see Supplementary Analyses and **Figure S2**).

Given that decisions involving trust rely on the interplay between brain regions and neural networks,^[15]^ we explored loneliness-related changes in functional connectivity by calculating generalized psychophysiological interaction (gPPI) analyses. The anatomically defined ROIs were used as seeds in seed-to-voxel analyses and trust-specific connectivity values (i.e. trust game > risk game) were compared between groups. Analyses revealed significant differences in the functional connectivity of the left AI with an occipitoparietal cluster including the cuneus and precuneus between LL and HL participants (−18, -76, 36, *k* = 163, *t*_(57)_ = 5.43, FWE-corrected *P* = 0.001 on cluster level; see **Figure 4**). Specifically, HL participants showed blunted functional connectivity of the left AI with this cluster during the trust game compared to LL participants (post-hoc *t*-test: *t*_(57)_ = -3.17, *P*_cor_ = 0.010, *d* = -0.83, 95 % CI of group difference: -0.45 to -0.10), whereas functional connectivity during the risk game did not significantly differ between groups (*t*_(57)_ = 1.59, *P*_cor_ = 0.472, 95 % CI of group difference: -0.04 to 0.34). Further post-hoc tests revealed increased functional connectivity during the trust game in LL participants (trust game versus risk game: *t*_(27)_ = 3.58, *P*_cor_ = 0.005, *d* = 0.49, 95 % CI of increase: 0.08 to 0.28), while connectivity during the trust game even decreased in HL participants (*t*_(30)_ = -4.16, *P*_cor_ = 0.001, *d* = -0.70, 95 % CI decrease: -0.36 to -0.12; for further analyses of connectivity, see Supplementary Analyses).

**Figure 4.**
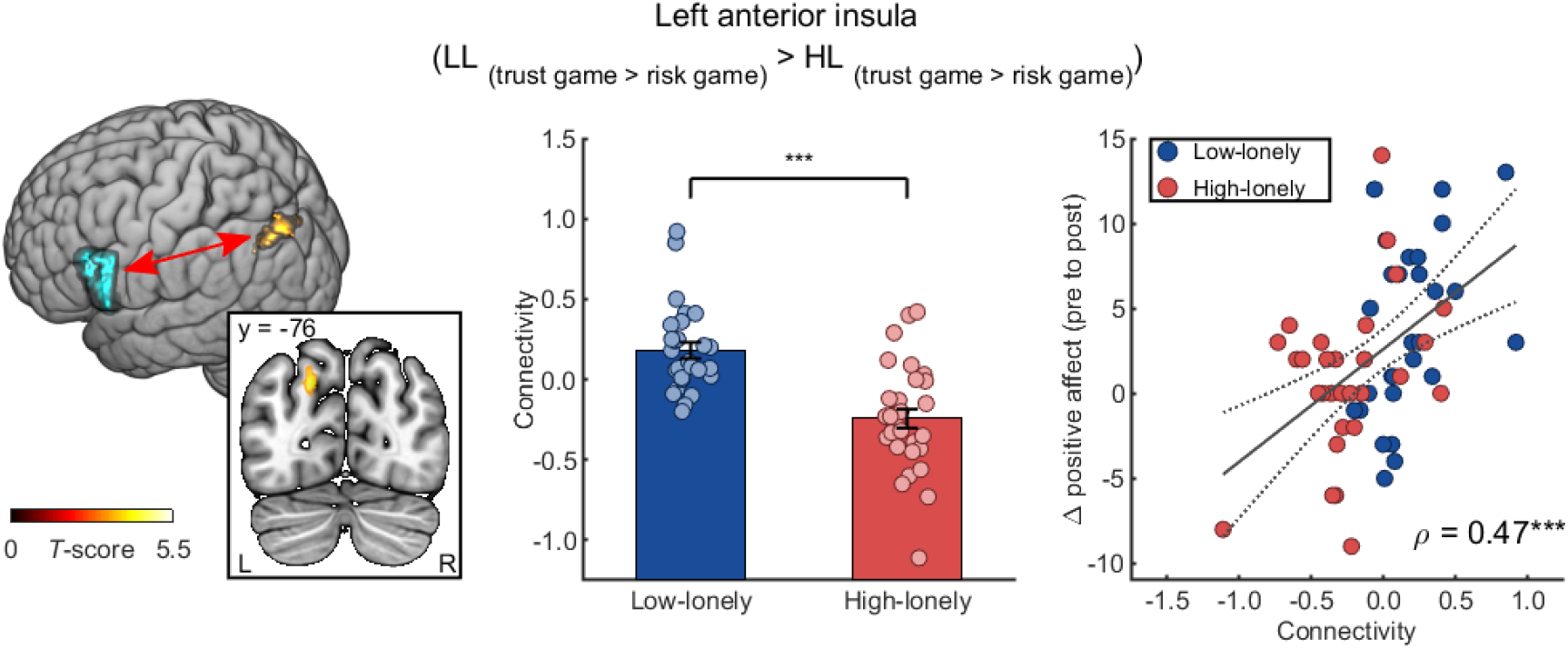
Reduced trust-associated connectivity of the anterior insula in high-lonely participants. High-lonely participants showed altered trust-associated connectivity of the left anterior insula with an occipitoparietal cluster including the cuneus and the precuneus. This connectivity of the left anterior insula during the trust game (compared to the risk game) positively correlated with the interaction-induced changes in positive affect across groups. The dashed line represents the 95%-confidence interval of the plotted regression line. All bars represent group means. Error bars indicate standard errors of the mean. Dots on bar plots are jittered for purposes of presentation. Abbreviations: HL, high-lonely; L, left; LL, low-lonely; R, right. *** P < 0.001.

Together, these results indicate that reduced interpersonal trust in HL participants might be based on an attenuated recruitment and functional connectivity of limbic regions and, more specifically, the AI during trust decisions. In a next step, we examined whether the observed differences in brain activity and connectivity were in fact associated with interpersonal trust measurements and with the attenuated responsiveness to the positive social interaction in loneliness.

### 2.4. Brain-behavior correlations

Our results confirmed that participants with less self-reported trust also showed less differentiated brain activity (left AI: ρ_(58)_ = 0.26, *P* = 0.047, 95 % CI: 0.004 to 0.48, see **Figure 3A**; right NAcc: ρ_(58)_ = 0.30, *P* = 0.020, 95 % CI: 0.05 to 0.52, see **Figure 3B**) and that greater trust-related increases in neural activity were associated with higher investments across conditions (left amygdala: ρ_(58)_ = 0.29, *P* = 0.028, 95 % CI: 0.03 to 0.51, see **Figure 3C**). Intriguingly, trust-specific connectivity of the left AI with occipitoparietal regions was positively associated with the social interaction-induced increase in positive mood (significant correlations with the increase in positive affect: ρ_(57)_ = 0.47, *P* = 0.0002, 95 % CI: 0.25 to 0.65, see **Figure 4**, and positive affect after the task: ρ_(57)_ = 0.30, *P* = 0.025, 95 % CI: 0.04 to 0.51, but not with baseline positive affect: ρ_(57)_ = -0.10, *P* = 0.439, 95 % CI: -0.35 to 0.16) indicating that impaired integration of trust-related information relates to diminished benefits of positive social interactions in HL participants. No further significant correlations were observed between neural and behavioral measurements and the oxytocinergic responsiveness to the positive social interaction.

Nevertheless, as the observed brain-behavior correlations might be driven by loneliness, we tested whether the reported correlations were also significant within each group. Analyses confirmed the positive association of AI connectivity with the positive affective responsiveness to the social interaction in LL participants (ρ_(27)_ = 0.47, *P* = 0.011, 95 % CI: 0.12 to 0.72). This correlation was not significant in HL participants (ρ_(29)_ = 0.25, *P* = 0.191, 95 % CI: -0.13 to 0.56). Moreover, the correlation of amygdala activity with the monetary investment during the trust and the risk game was found for the HL participants (ρ_(30)_ = 0.43, *P* = 0.016, 95 % CI: 0.09 to 0.68) but was absent in LL participants (ρ_(27)_ = 0.15, *P* = 0.458, 95 % CI: -0.24 to 0.49). None of the other reported correlations reached significance within the groups.

### 2.5. Loneliness, subclinical psychiatric symptoms, and sex differences

HL participants were characterized by heightened depressive and anxiety symptoms, childhood maltreatment, and worse sleep quality (all *P*s < 0.020; see **Table S4**). Furthermore, HL participants reported smaller and less diverse social networks (all *P*s < 0.002; see **Table S4**). To assess whether the observed associations of loneliness with behavioral, neuroendocrine, and neural alterations are mediated by these psychiatric characteristics of HL participants, we conducted mediation analyses with depressive and anxiety symptoms and childhood maltreatment scores as mediator variables and group as independent variable. None of the reported group effects was confounded by any of the tested mediators (all 95 % CIs of mediation effects overlapped with zero) except for the reduced AUC_I_ in salivary oxytocin levels after completion of the positive social interaction task which was mediated by depressive symptomatology (β = 0.22, SE = 0.11, 95 % CI: 0.04 to 0.47).

Notably, the sex of the participants did not significantly influence the strength of the reported associations of loneliness with brain activity or connectivity: No interactions of group with sex were observed for trust-associated AI activity (group-by-sex interaction: *B* = 0.12, *t*_(55)_ = 1.04, *P* = 0.302, 95 % CI: -0.11 to 0.35) or connectivity (*B* = -0.11, *t*_(55)_ = -0.69, *P* = 0.496, 95 % CI: -0.42 to 0.21) or for trust-associated amygdala (*B* = -0.02, *t*_(55)_ = -0.19, *P* = 0.847, 95 % CI: -0.25 to 0.21), NAcc (*B* = -0.16, *t*_(55)_ = -1.53, *P* = 0.131, 95 % CI: -0.37 to 0.05), or mPFC activity (*B* = 0.20, *t*_(55)_ = 0.65, *P* = 0.522, 95 % CI: -0.42 to 0.83). Likewise, the sex of the participants did not significantly influence other findings, although the association between loneliness and general trust appears to be more pronounced in men than women (group-by-sex interaction: *B* = -0.43, *t*_(78)_ = -1.82, *P* = 0.073, 95 % CI: -0.90 to 0.04; group effects in female participants: *B* = -0.34, *P* = 0.048, 95 % CI = -0.67 to -0.002; male participants: *B* = -0.77, *P* < 0.001, 95 % CI = -1.10 to -0.43; all further interactions of group with sex: *P*s > 0.130). For further moderation analyses, see Supplementary Analyses.

## 3. Discussion

Our study sought to investigate a trust-based mechanism underlying the attenuated reactivity to positive social interactions in a pre-stratified sample of HL and LL participants. As hypothesized, HL individuals exhibited reduced affective responses to the positive social interaction and reported less interpersonal trust. Moreover, during initial trust decisions, blunted AI activity in HL participants was consistently found across frequentist and Bayesian analyses and was accompanied by reduced functional connectivity of the AI to an occipitoparietal cluster, including the cuneus and precuneus, which correlated with attenuated affective reactivity to the positive social interaction. Frequentist analyses further indicate diminished trust-associated brain activity in the amygdala and NAcc, while Bayesian analyses provide strong evidence for blunted mPFC activity in HL participants. Further explorative analyses revealed attenuated oxytocinergic responsiveness to the positive discussion in HL participants and that HL participants were rated to be less trustworthy by the unfamiliar experimenter. Notably, although the HL sample was characterized by heightened psychiatric symptomatology, neither depression or social anxiety scores nor reported childhood maltreatment mediated the observed neural group differences.

Our results confirmed the findings of previous studies reporting reduced responsiveness to positive social interactions in lonely individuals:^[12,13]^ while the LL sample showed the expected increase in positive affect and salivary oxytocin concentrations, these responses were significantly diminished in HL participants. Furthermore, consistent with previous observations,^[28]^ HL participants preferred a greater interpersonal distance. Although this greater interpersonal distance might also reflect safety behavior due to the weakened immune system in lonely individuals (^[10]^ also see SI), our results strongly support theoretical framework suggesting that negative biases have a detrimental effect on social interactions in loneliness,^[11]^ as impaired interpersonal trust significantly correlated with a preference for larger interpersonal distance and reduced positive mood after the positive social interaction.

Mechanistically, our findings might indicate that impaired trust evaluations could be rooted in attenuated limbic reactivity. Our observation of reduced trust-associated activation in the AI is consistent with recent studies highlighting the AI as a key region contributing to trust decisions specifically during the single-round trust game.^[29]^ The AI encodes the trustworthiness of faces^[30]^ and is fundamentally involved in integrating interoceptive information from limbic regions including the amygdala and the NAcc. Interestingly, the AI initializes the processing of salient information in the prefrontal cortex, encodes the incentive value of stimuli,^[31]^ and changes in the glucose metabolism of the AI positively correlate with changes in social interest.^[32]^ Thus, the reduced trust-related AI activity might indicate a compromised integration of amygdalar and striatal trust signals that might contribute to the overall nihilistic feeling that nobody can be trusted. Moreover, HL participants showed an altered interplay of the AI with a brain cluster including parts of the precuneus. The precuneus is a central hub of the default mode network and contributes to self-referential operations including self-consciousness and the mental representation of the self.^[33]^ Importantly, the functional connectivity of the precuneus with the AI during rest has been previously found to predict trust and reciprocity in non-lonely individuals.^[34,35]^ During positive social interactions, the precuneus might contribute to the continuously updated representation of a positive self-image that could reinforce the reward value of social interactions.^[36]^ In fact, connectivity of the AI with the precuneus correlated with the beneficial effects of our positive social interaction.

Furthermore, our results indicate a diminished recruitment of the amygdala, the NAcc, and the mPFC during trust-processing in lonely individuals. The mPFC has been previously implicated in loneliness,^[14,37]^ but our findings have to be interpreted with caution as the reduced trust-associated activity in HL participants could not be replicated across different analytic approaches. Like the AI, the mPFC is known to interact with various limbic regions, encode the expected value of stimuli,^[38,39]^ evaluate trait characteristics of others,^[40]^ and predict trusting behavior.^[34,41]^ The observed attenuated mPFC activity during the trust game might thus reflect a reduced utility of social stimuli, as lonely individuals potentially prefer safety behavior irrespective of the trustworthiness of the partner.^[42]^ In addition, the reduced mPFC activity might be linked to the attenuated recruitment of the amygdala and the NAcc.^[38,42]^

The amygdala is crucially involved in the processing of social information, such as the trustworthiness or ambiguity of social stimuli, and previous lesion studies provide strong evidence that an intact amygdala is necessary for developing appropriate interpersonal trust.^[30,43]^ Notably, like the AI, the amygdala encodes not only the negative valence of stimuli but also signals highly untrustworthy and trustworthy faces.^[30]^ Together with the association of reduced amygdala reactivity and lower monetary investment across conditions, our results could indicate that HL individuals might be less able to reliably evaluate the trustworthiness of strangers. This way, reduced amygdala sensitivity for trustworthiness evaluations might be a reinforcing mechanism for a default distrust mode as safety behavior in loneliness.

Moreover, intact amygdala projections to the NAcc are important to guide action selection in situations involving reward uncertainty^[44]^ and the NAcc showed diminished activity during trust decisions in HL individuals. The NAcc consistently responds to trust decisions during the multi-round trust game,^[29]^ but since we implemented a single-round version of the trust game, the striatal hypoactivation might reflect a general attitude of reduced trust towards strangers rather than previous learning experiences with the individual trustees. Nevertheless, our results might also indicate a reduced reward value of social stimuli in loneliness *per se*^[12]^ irrespective of the expected outcome during the trust game.

Notably, HL participants did not differ from the LL sample in trust-related activity of the TPJ, known to play a crucial role in inferring the mental state and temporary goals of other persons.^[17,45]^ As such, our findings point to a compromised integration of interoceptive trust signals and mental self-representation mediated by the functional interplay between the AI and precuneus as well as an impaired processing of trustworthiness and stimulus utility in the amygdala, NAcc, and mPFC, rather than altered inferences about the mental states of others as primarily processed in the TPJ.

Of note, diminished reactivity to social interactions in HL individuals was not limited to self-reported mood but also evident in significantly lowered endogenous oxytocin responsiveness. Oxytocin is crucially involved in human affiliation and trust^[23-25]^ and recent studies have highlighted the potential of intranasally administered oxytocin to increase interpersonal trust behavior in participants with a low disposition to trust.^[46,47]^ We have previously found that social synchrony, i.e. the temporal coordination of social behavior and physiological processes among individuals, evokes heightened endogenous oxytocin release, which predicts interactive reciprocity^[48]^ and that intranasal administration of oxytocin increases synchrony during dance.^[49]^ Social synchrony is essential for human bonding and has been associated with positive affect and prosocial behavior.^[50]^ While the electrodermal and heart rate measurements demonstrate a normal arousal response to the positive social interaction, HL individuals not only reported less interpersonal trust but they were also rated as less trustworthy and the blinded experimenter was able to recognize them better than by chance. Thus, our findings support previous reports about the social transmission of loneliness^[27]^ and suggest that the impaired trust evaluation may hamper social synchrony, which in turn can explain the lower perceived trustworthiness of HL individuals.^[51]^ Along these lines, dysfunctional social interactions in loneliness may result from a reciprocally-reinforced bias in trust behavior.

The current study has several limitations. The cross-sectional design of the current study does not allow causal inferences about the relationship between interpersonal trust, loneliness, or the beneficial effects of positive social interactions. Although preliminary evidence supports the notion of reduced interpersonal trust as a risk factor for rather than a consequence of loneliness,^[11]^ future longitudinal studies are required to directly test the causality of this model. Likewise, experimental studies using neurofeedback training, human lesion models, or transient lesions via non-invasive brain stimulation are needed to prove the causal involvement of the observed trust-associated neurocircuit in loneliness. Furthermore, interpersonal synchrony needs to be characterized in naturalistic settings with two HL participants and mixed dyads of HL and LL individuals. Notably, although moderation analyses did not reveal significant interactions of loneliness with the participants’ sex, this does not exclude the possibility of sex differences in other loneliness-related domains or population-based measurements. For instance, previous studies found sex-specific associations of loneliness with brain structure and resting state functional connectivity using the UK biobank population.^[14,52,53]^

## 4. Conclusion

Collectively, our results indicate compromised integration of trust-related information as a potential reciprocally-reinforced mechanism that might contribute to dysfunctional social interactions in loneliness, thereby reducing the motivation to reconnect and promoting avoidance behavior. Neurobiologically-informed interventions with cognitive bias modification procedures should target the self-reinforcing loop of distrust to improve the beneficial reactivity to positive social interactions and alleviate the debilitating health consequences of perceived social isolation.

## 5. Methods

### Participants and study design

To investigate the impact of current loneliness on interpersonal trust, the reactivity to positive human interactions, and its underlying neurobiological mechanisms, we used a quasi-experimental design with a sample of pre-stratified healthy volunteers scoring high (≥ 50, i.e. at least one standard deviation above the mean score of students, cf.^[54]^) or low (≤ 25, i.e. at least one standard deviation below the mean) on the revised UCLA Loneliness Scale (UCLA-L).^[54]^ For recruitment, an online survey assessing the UCLA-L score was disseminated by means of online advertisement and public postings. A total of 410 participants out of 3678 subjects who filled out the UCLA-L scale met the inclusion criteria (see Supplementary Methods) and out of these 410 participants, 91 subjects agreed to participate and were invited to a screening session. Nine participants were excluded after the screening session since they were not eligible for enrolment, resulting in a final sample of 42 HL (female *n* = 21) and 40 LL participants (female *n* = 20) in accordance with the planned sample size of 80 participants (for details of the a-priori power analysis, see Supplementary Methods). Groups were matched for age (HL mean age ± SD: 26.55 ± 6.80 years, LL: 27.13 ± 8.18 years; *t*_(80)_ = 0.35, *P* > 0.05, 95 % CI of group difference: -3.88 to 2.72 years) and sex and did not differ regarding sociodemographic factors (all *P*s > 0.05; see **Table S1**). All participants provided written informed consent and received monetary compensation for participation. The study was approved by the local ethics committee of the Medical Faculty of the University of Bonn, Germany (study number 016/18), and carried out in accordance with the latest revision of the Declaration of Helsinki. Data analysis was preregistered prior to conducting any analyses (https://osf.io/x47ke; results regarding the preregistered hypothesis #3 will be published elsewhere).

### Psychological variables

Participants completed questionnaires measuring interpersonal trust, the social network size and diversity, and sleep quality. As loneliness is often associated with psychiatric symptomatology, we also assessed depressive symptoms, social anxiety, and childhood maltreatment. For details, see Supplementary Methods.

### Trust game

We implemented an adapted version of an established trust game (cf.^[21]^). Briefly, two players, the investor and the trustee, start each round with an endowment of 10 €. The investor chooses the amount of money he/she wants to invest in an unknown trustee. The invested money is tripled and added to the trustee’s account. The trustee can keep all of the money for him/herself or share the money with the investor so that both players end with the same amount of money (10 € plus the invested amount). Decisions of the participants in the role of the trustee were collected for all possible investments during the screening session (see also Supplementary Analyses). Participants were informed that they would play the trust game in the role of the investor against other participants of the study (as trustees) and that their own payment depended on a randomly chosen trial (100 % of the final endowment after consideration of the trustee’s decision was paid). During fMRI, participants then played the trust game as investor without receiving feedback about the pre-recorded decisions of the trustees to explore the impact of loneliness on rapid trustworthiness decisions during initial encounters. In a control condition, participants played a risk game in which they invested money in a computer (which would randomly decide whether the money would be shared).

As choice options and possible outcomes were exactly the same during the trust game and the risk game, the conditions differed only with respect to the social risk of betrayal when playing with a human counterpart. Thus, when analyzing trust-related decisions and associated brain activity, it is crucial to validate whether participants believed they were playing against real persons as no differences should be observed otherwise. We therefore asked participants both verbally and via questionnaire whether they believed our instructions. For details, see Supplementary Methods.

### Positive social interaction paradigm

After completion of the fMRI scan, participants moved to the testing room, which was prepared for the positive social interaction paradigm. The task consisted of a semi-structured 10-minute conversation between the participant and a same-sex unfamiliar experimenter. Participants were told to talk about (1) plans for a fictive lottery win, (2) positive childhood memories, and (3) hobbies and interests. High-quality photographs presenting examples for activities (e.g. traveling around the world or buying a sports car) were used to facilitate the start of the conversation. Participants and the experimenter tried to find similarities in the discussed topics. Importantly, the experimenter was blinded regarding the group assignment of the participant (HL vs. LL) and unknown to the participants prior to the fMRI session in all cases. Participants self-reported mood and affect before and after completing the positive social interaction paradigm (see Supplementary Methods). After finishing the task, both the participant and experimenter, rated the valence of the discussion as well as trustworthiness and likeability of each other using VAS. The experimenter further estimated the experimental group of the participant (HL vs. LL) to examine whether loneliness might be detected by others after the positive interaction.

Baseline electrodermal activity (EDA) and an electrocardiogram (ECG) were collected for five minutes prior to the positive social interaction paradigm and throughout the entire social interaction. Finally, saliva samples were collected before, immediately after the social interaction paradigm, and 15 minutes after completion of the positive social interaction task to obtain salivary oxytocin, cortisol, and IgA levels in addition to baseline immune parameters and oxytocin levels in blood (see Supplementary Methods).

### Interpersonal distance paradigm

The interpersonal distance as an indirect index of trust towards strangers was measured by an adapted version of an established stop-distance paradigm (cf.^[22]^). Participants moved toward an unfamiliar experimenter (the same experimenter who conducted the positive social interaction) from a start distance of 2 m and stopped at their ideal distance. In a second trial, participants were instructed to stop at a distance at which they felt slightly uncomfortable. The start and final chin-to-chin distance were measured with a digital laser measurer (error: ± 0.003 m). Both conditions were measured before and after the positive social interaction.

### fMRI data analysis

fMRI data were acquired with a 3T Siemens TRIO MRI system (Siemens AG, Erlangen, Germany) using a T2*-weighted echoplanar (EPI) sequence and preprocessed and analyzed using standard procedures in SPM12 (Wellcome Trust Center for Neuroimaging, London, UK; http://www.fil.ion.ucl.ac.uk/spm) implemented in Matlab (The MathWorks Inc., Natick, MA) (see Supplementary Methods). A two-stage approach based on the general linear model implemented in SPM12 was used for statistical analyses. On the first level, participants’ individual data were modelled using a fixed-effects model. Onsets and durations of the experimental conditions were modelled by a stick function convolved with a hemodynamic response function (HRF). Movement parameters were included in the design matrix as confounds. In line with previous research investigating trust (cf.^[21]^) and according to our hypothesis of altered trust-associated brain activity in HL participants, we contrasted individual brain activity during the trust game with the risk game (trust game > risk game) as main contrast of interest on the first level and compared groups by using two-sample *t*-tests on the second level (HL _trust game > risk game_ > LL _trust game > risk game_, LL _trust game > risk game_ > HL _trust game > risk game_; for further details and analyses, see Supplementary Methods). The main analyses of fMRI data focused on independently defined brain regions (ROIs) known to be involved in motivational, affective and cognitive processes during the trust game consisting of the bilateral amygdala, AI, mPFC, and TPJ^[15]^ (see Supplementary Methods and **Figure S3**). We further included the NAcc as ROI associated with reward anticipation during the trust decision stage because the NAcc has been found to show altered activity as a function of loneliness.^[12]^ *P* values smaller than 0.05 after FWE-correction based on the size of the ROI (i.e. small volume correction) were considered significant. Whole-brain analyses were calculated across groups for task validation. Parameter estimates of significant contrasts were extracted using marsbar (http://marsbar.sourceforge.net) and further analyzed to disentangle interactions by calculating Bonferroni-corrected post-hoc *t*-tests. To account for the liberal threshold of small volume corrections in our ROI analyses, we probed the robustness of observed group differences in trust-associated brain activity by conducting Bayesian inference analyses as implemented in SPM (cf.^[55]^). Results were thresholded with the following criteria: a log odds Bayes factor threshold of logBF ≥ 3 (strong evidence,^[56]^) for at least small group effects (i.e. an effect size threshold of 0.2). A gPPI analysis was conducted using the CONN toolbox 19.b (www.nitrc.org/projects/conn, RRID:SCR_009550) and the same statistical model as outlined above (for details, see Supplementary Methods). Those ROIs that showed significant effects during the fMRI trust game (i.e. left amygdala, left AI, right NAcc) were used as seed regions in planned seed-to-voxel analyses, while all other ROIs were used as seed regions in additional exploratory seed-to-voxel analyses (see Supplementary Analyses). For each participant, interaction terms of the psychological factor (effects of task conditions convolved with a canonical HRF) and the physiological factor (seed ROI BOLD time series) were computed on the first level. Bivariate regression measures were used to provide the relative measure of connectivity compared to the implicit baseline (defined by the zero values of the interaction term). On the second level, we compared trust-specific connectivity values between groups using 2×2 mixed ANOVA interactions (HL _trust game > risk game_ > LL _trust game > risk game_; LL _trust game > risk game_ > HL _trust game > risk game_) to test the hypothesis of altered connectivity in loneliness. Results were thresholded at an FWE-corrected *P*-value < 0.05 after an initial cluster-forming height threshold of *P* < 0.001. Beta weights of significant effects of interest were extracted and further analyzed by calculating Bonferroni-corrected post-hoc *t*-tests.

### Behavioral and questionnaire data analysis

Statistical analyses were performed using SPSS 24 (IBM Corp., Armonk, NY). Questionnaire data were compared between groups using two-sample *t*-tests and chi-square tests. All behavioral data were analyzed using mixed-design ANOVAs and Bonferroni-corrected post-hoc *t*-tests. If the assumption of sphericity was significantly violated as assessed by Mauchly’s tests, Greenhouse-Geisser corrections were applied. The sociality condition of the trust game served as within-subject factor (trust game vs. risk game), while group constituted the between-subject factor (HL vs. LL). The hypothesized group differences in the response to the positive social interaction paradigm (self-reported affect and mood) and the interpersonal distance task (separated for comfortable and uncomfortable distance) were analyzed with time (before vs. after social interaction) as within-subject factor and group as between-subject factor. Analyses of the trust game excluded participants who did not believe the instructions as stated verbally or during the exit questionnaire (*n* = 8 HL, *n* = 9 LL). For analyses of the positive social interaction paradigm, participants who were not fluent in German were excluded (*n* = 3 HL). Chi-square tests were used to calculate whether the estimation of the experimental group (HL vs. LL) by the experimenter differed significantly from chance.

### Psychophysiology and neuroendocrinology analysis

The SCL and heart rate (BPM) were analyzed using mixed ANOVAs including the within-subject factor time (baseline vs. social interaction) and the between-subject factor group (HL vs. LL). The difference between the duration of the baseline acquisition and the duration of the social interaction task was included as covariate to control for changes in psychophysiology related to differences in data acquisition times.

Baseline differences in the salivary and plasma oxytocin levels, salivary cortisol and IgA concentrations, and in blood parameters (serum C-reactive protein, interleukin-6, 25-hydroxyvitamin D, oxytocin, and cell count parameters) were compared between groups using two-sample *t*-tests. The AUC_I_ (see Supplementary Methods) was calculated for salivary oxytocin, cortisol, and IgA levels, tested against zero to examine the responsiveness to the positive social interaction across groups and compared between groups, again using two-sample *t*-tests.

### Correlation, mediation, and moderation analyses

To examine the hypothesis that altered brain activity and connectivity in HL participants relate to the observed behavioral group differences, we correlated parameter estimates of trust-specific brain activity and connectivity with the behavioral variables that were associated with loneliness (for details, see Supplementary Methods). To further explore the relationship of interpersonal trust with behavioral data, we also correlated the self-reported interpersonal trust with those variables. Furthermore, correlation analyses were calculated separately for each group.

To examine whether observed group effects (main effects of group or interactions with group) might be driven by psychiatric symptomatology, we calculated mediation analyses and tested for indirect effects of loneliness via psychiatric symptomatology. Thus, we examined whether the observed effects of loneliness might be partially or fully based on the psychiatric symptoms associated with loneliness.

In addition, to expand our understanding of the interplay of loneliness and psychiatric symptomatology, moderation analyses were conducted to investigate potential interaction effects. This way, we were able to test whether psychiatric symptomatology might potentiate observed effects associated with loneliness (i.e. stronger effect of loneliness in participants with higher psychiatric symptoms) or reduce the impact of loneliness (i.e. less effect of loneliness in participants with higher psychiatric symptoms). Likewise, we conducted moderation analyses with the sex of the participants as moderator variable to examine whether the effects of loneliness differed between sexes. For details, see Supplementary Methods.

[The behavioral data that support the findings of the current study are openly available in the repository of the Open Science Foundation at https://osf.io/p6jxk/. The unthresholded statistical maps of the fMRI results can be accessed at https://neurovault.org/collections/BMDUCOHK/.]

## Acknowledgments

J.L. and D.S. designed the experiment; J.L., T.E., and E.K. conducted the experiments; J.L., E.K., and D.S. analyzed the data. All authors wrote the manuscript. All authors read and approved the manuscript in its current version. The authors declare no competing interests. The authors thank Paul Jung for programming assistance and Alexandra Goertzen-Patin for proofreading the manuscript. S.G.S-T, R.H., and D.S. are supported by a German-Israel Foundation for Scientific Research and Development grant (GIF, I-1428-105.4/2017).

