## Supporting Information for "Loneliness and the social brain: how perceived social isolation impairs human interactions"

### Supplementary Methods

#### Inclusion and exclusion criteria

Participants were pre-stratified and assigned to the high- (HL) or low-lonely (LL) group using the UCLA Loneliness Scale (UCLA-L).[1,2] A total of 410 participants out of 3678 subjects who filled out the UCLA-L scale met the following inclusion criteria: UCLA-L score ≥ 50 or ≤ 25, resp., no current physical or psychiatric illness, right-handed, eligibility for MRI scanning, aged 18 to 65, consent to be contacted. Exclusion criteria further consisted of current psychotherapy, medication (except for hormonal contraceptives, thyroid medicines, or asthma inhalers), or illicit drug use in the previous four weeks. The Mini-International Neuropsychiatric Interview (MINI)[3] was conducted during the screening session to assess possible psychological disorders.

#### Power analysis

We used G*Power 3[4] to conduct an a-priori power analysis for the project to determine the planned sample size. To reliably replicate the previously observed effect of perceived social isolation on the neural processing of social stimuli[5] (with $\alpha = 0.05$ and power = 0.99), at least 71 participants had to be tested in a between-subject design. To account for possible drop-outs, we planned to test at least 80 participants.

#### Questionnaires

Participants completed questionnaires measuring the social network size and diversity[6] and sleep quality[7] in addition to sociodemographic characteristics including monthly salary ("0 to 500 €", "501 to 1,000 €", "1,001 to 1,5000 €", "1,501 to 2,000 €", "2,001 to 2,500 €", "2,501 to 3,000 €", "more than 3,000 €"). Depressive symptoms, social anxiety, and childhood maltreatment were assessed using the Beck Depression Inventory-II (BDI)[8], the Liebowitz Social Anxiety Scale (LSAS)[9], and the Childhood Trauma Questionnaire (CTQ)[10]. Interpersonal trust as trait variable was measured with the General Trust Scale [11]. Mood was assessed by the Positive and Negative Affect Scales (PANAS)[12], the Profile of Mood States (POMS) questionnaire[13], and visual analogue scales (VAS, ranged 0 to 100 and measuring loneliness, happiness, sadness, anger, anxiety, and overall arousal) before and after completing the positive social interaction paradigm.

**fMRI trust game implementation**

To increase the credibility of the instructions that all participants played the trust game against other, randomly chosen participants, a photograph of each participant was taken during the screening session and participants were told that their photograph may be used during the fMRI session of other participants. Furthermore, participants were informed that their own decisions as trustee could influence the payment for other participants playing the trust game as investor during fMRI and participants knew that their own payment depended on a randomly chosen trial as trustee (10 % of the kept amount was paid) as well. While collecting the trustee's decisions, visual aids were given to ensure the participants' understanding of the effect of their decisions on the final credit of both players.

During fMRI, participants first completed training sessions of the trust game inside the scanner to get used to the design of the task and to respond within the fixed time interval of 7 s. Participants could repeat the training as often as needed to feel comfortable starting the task. Participants were reminded that they would receive the money of a randomly chosen trial depending on their investment and the pre-recorded decision of the trustee. A photograph of the putative trustee was shown in each trial. In reality, pictures of the Chicago Face Database[14] were used as photographs for reasons of standardization and to guarantee the privacy of the participants. We selected pictures of Caucasian persons aged 20 to 30 years to ensure the comparability with our sample consisting of mostly German students and thus to increase the credibility of the cover story. Neutral expression photographs consisted of 15 Caucasian female actors (i.e. WF-003, WF-006, WF-008, WF-009, WF-012, WF-017, WF-021, WF-027, WF-028, WF-035, WF-207, WF-214. WF-218, WF-219, WF-230) and 15 Caucasian male actors (i.e. WM-003, WM-006, WM-009, WM-015, WM-024, WM-203, WM-205, WM-208, WM-210, WM-212, WM-213, WM-214, WM-245, WM-247, WM-253). Female and male actors did not differ regarding estimated age, attractiveness, and trustworthiness as indicated by the norming data (all $P$s > 0.193). Computer stimuli were matched in width and height to social stimuli and depicted computers in front of a white background.

In each trial, pictures were presented on a black background using the software Presentation 17 (Neurobehavioral Systems, Albany, CA, USA). The order of conditions (trust game, risk game, age human, age computer) was quasi randomized (no more than two trials of the same condition in a row). Participants could choose their responses using an MRI-compatible response grip system (NordicNeuroLab AS, Bergen, Norway). Two buttons could be used with the index fingers to move a cursor left and right on a white scale below the picture. Participants confirmed their responses by pressing two buttons with their thumbs. After confirmation, the cursor changed to green and was not movable anymore. For each position, the current value (invested money or estimated age) was presented in the middle of the screen. In each condition, the scale included a range of 11 values (trust and risk game: 0 to 10 €, age human: 20 to 30 years, age computer: 0 to 50 years in 5-year increments). The direction of the scale (i.e. 0 to 10 € or 10 to 0 €) was counterbalanced across participants. For optimization of the design, the easy-optimize-x software was used [https://www.bobspunt.com/easy-optimize-x/; 4 conditions with 30 trials each, maximum block size: 2, trial duration: 7 s, mean inter stimulus interval (ISI): 5 s (range: 4-6 s), ISI distribution: Rayleigh, time before first and after last trial: 0 s, the best 85 designs were saved out of 50 generations (*N* designs per generation: 2000), max time to run: 1440 min, 3 contrasts of interest: (1 -1 0 0; 0 0 1 -1; 1 -1 -1 1)]. One design was calculated for each participant. We asked participants both verbally ("Did you believe that you played against other participants?") and via questionnaire ("Did you believe that the photographs during the fMRI trust game presented other participants?") whether they believed our instructions after being informed about the cover story.

**fMRI data acquisition and preprocessing**

Functional data of the trust game were acquired with a 3T Siemens TRIO MRI system (Siemens AG, Erlangen, Germany) with a Siemens 32-channel head coil and obtained using a T2*-weighted echoplanar (EPI) sequence [TR = 2690 ms, echo time (TE) = 30 ms, ascending slicing, matrix size: 96 x 96, voxel size: 2 x 2 x 3 $mm^3$, slice thickness = 3.0 mm, distance factor = 10 %, field of view (FoV) = 192 mm, flip angle 90°, 41 axial slices]. High-resolution T1-weighted structural images were collected on the same scanner (TR = 1660 ms, TE = 2.54 ms, matrix size: 320 x 320, voxel size: 0.8 x 0.8 x 0.8 $mm^3$, slice thickness = 0.8 mm, FoV = 256 mm, flip angle = 9°, 208 sagittal slices). To control for inhomogeneity of the magnetic field, fieldmaps were obtained for each T2*-weighted EPI sequence [TR = 392 ms, TE (1) = 4.92, TE (2) = 7.38, matrix size: 64 x 64, voxel size: 3 x 3 x 3, slice thickness = 3.0 mm, distance factor = 10 %, FoV = 192 mm, flip angle 60°, 37 axial slices]. Participants with excessive head movements (> 4 mm/° in any direction, *n* = 3 HL, *n* = 3 LL) or anatomical abnormalities (*n* = 1 LL) were excluded from fMRI analyses. The first five volumes of each functional time series were discarded to allow for T1 equilibration. Functional images were corrected for head movements between scans by an affine registration. Images were initially realigned to the first image of the time series before being re-realigned to the mean of all images. To correct for signal distortion based on B0-field inhomogeneity, the images were unwarped by applying the voxel displacement map (VDM file) to the EPI time series (Realign & Unwarp). Normalization parameters were determined by segmentation and non-linear warping of the structural scan to reference tissue probability maps in Montreal Neurological Institute (MNI) space. Normalization parameters were then applied to all functional images, which were resampled at 2 x 2 x 2 $mm^3$ voxel size. For spatial smoothing, a 6-mm full width at half maximum (FWHM) Gaussian kernel was used. Raw time series were detrended using a high-pass filter (cut-off period 128 s). Furthermore, preprocessing for functional connectivity analyses included a denoising pipeline. Following the recommendations, outlier scans were detected by the integrated artefact detection toolbox-based identification using conservative settings (i.e. thresholds of 0.5 mm frame wise displacement and 3 SD above global BOLD signal changes were used). The default denoising pipeline implemented a linear regression of confounding effects of the first five principal noise components from white matter and cerebrospinal fluid template masks, 12 motion parameters, scrubbing, and constant task-related effects. A high-pass filter of 0.008 Hz was applied to minimize the effects of physiological and motion related noise.

### Exploratory fMRI data analyses and regions of interest

To further control for the social content of the stimuli, we included an additional condition in which participants had to estimate the age of the presented social and computer stimuli. For the first level analyses, one regressor for each of the four conditions (‘trust game’, ‘risk game’, ‘age human’, ‘age computer’) was included in the design matrix in addition to the six movement regressors and the modelled baseline. In addition to the main contrasts of interest focusing on the comparison of the trust game with the risk game, we explored whether trust-related brain activity differed between groups when contrasting the trust game with this social but not non-economic control condition [i.e. $HL_{\text{trust game > age human}} > LL_{\text{trust game > age human}}$, $LL_{\text{trust game > age human}} > HL_{\text{trust game > age human}}$, $HL_{\text{age human > age computer}} > LL_{\text{age human > age computer}}$, $LL_{\text{age human > age computer}} > HL_{\text{age human > age computer}}$, $HL_{\text{(trust game > risk game) > (age human > age computer)}} > LL_{\text{(trust game > risk game) > (age human > age computer)}}$, and $LL_{\text{(trust game > risk game) > (age human > age computer)}} > HL_{\text{(trust game > risk game) > (age human > age computer)}}$].

The regions of interest (ROIs) included the amygdala, the anterior insula (AI), the medial prefrontal cortex (mPFC), and the nucleus accumbens (NAcc) and were anatomically defined according to the Wake Forest University PickAtlas.[15,16] Specifically, the amygdala and the AI were defined according to the aal atlas. To create the structural ROI of the AI, a caudal boundary of y = 8 was applied to the original insula mask.[17,18] The anatomical masks of the bilateral medial frontal gyrus, as defined in the TD labels atlas, and the NAcc, as included in the IBASPM 71 atlas, were used as mPFC and NAcc, respectively. Furthermore, the temporoparietal junction (TPJ) was in,dependently created using association test maps from neurosynth (https://neurosynth.org/; search term: temporoparietal junction) with peak voxels in the left (-50, -56, 22; cluster size: 704 voxels) and right (54, -56, 20; cluster size: 1439 voxels) TPJ. All ROIs are displayed in Figure S3.

**Neuroendocrine parameter extraction and analysis**

Saliva samples were collected using commercial sampling devices (Salivettes, Sarstedt, Germany) and immediately cooled after collection. Saliva samples were centrifuged at 4,000 rpm for 2 minutes and stored at -80°C until assayed. Additionally, at the beginning of the fMRI session, venous ethylenediaminetetraacetic acid (EDTA) whole blood samples were collected for the assessment of levels of erythrocytes, leucocytes, thrombocytes, lymphocytes, basophile granulocytes, neutrophil granulocytes, eosinophil granulocytes, and monocytes (cells as absolute and percentage count). Blood was immediately transported to the central laboratory of the University Hospital Bonn. In all samples, analysis was performed within 15 minutes of arrival at the central laboratory. A further plasma sample was collected with commercial sampling devices (Vacuette, Greiner Bio-One International, Austria) containing EDTA and aprotinin for oxytocin concentrations assessment, centrifuged at 4,000 rpm for 20 minutes and stored at -80°C until assayed.

A highly sensitive and specific radioimmunoassay (detection limit of 0.1 - 0.5 pg, depending on the age of the tracer) was used to extract and quantify salivary and plasma oxytocin concentrations (RIAgnosis, Munich, Germany)[19]. Salivary cortisol and serum interleukin-6 (IL-6) levels were analyzed with the electro chemiluminescence assays (ECLIA) for Cobas™ e801 analyzer (Roche Diagnostics GmBH, Mannheim, Germany; detection limit of 1.5 - 1,750 nmol/L for cortisol and 1.5 – 5,000 pg/mL for IL-6) and measurements for immunoglobulin A (IgA) in saliva were performed with particle-enhanced immunological turbidity assay for Cobas™ c502 analyzer (Roche Diagnostics; detection limit: 0.1 - 6 g/L). Measurements for serum C-reactive protein

(CRP) were performed with an immunological turbidity assay for Cobas c702™ analyzer (Roche Diagnostics; detection limit of 0.6 – 350 mg/L). An automated CE-IVD workflow solution was used for determination of serum 25-OH-vitamin D3 levels. It included validated, automated sample preparation with the MassSTAR system (Hamilton Germany GmbH, Martinsried, Germany) and liquid chromatography-tandem mass spectrometry (LC-MS/MS) with Q-TRAP 4500MD (AB SCIEX Germany GmbH, Darmstadt, Germany) using the MassChrom™ 25-OH-vitamin D3/D2 assay (Chromsystems, Munich, Germany; detection limit of 1.0 – 250 µg/l) according to the manufacturer's instructions. Intra- and inter-assay coefficients of variability were < 20 % for all assays. All samples to be compared were assayed in the same batch, i.e. under intra-assay conditions. Further blood parameters were analyzed with Sysmex XN1000™ (Sysmex, Norderstedt, Germany). Erythrocytes and thrombocytes were measured with impedance technique. In the Sysmex WDF channel, lymphocytes, monocytes, neutrophil granulocytes, and eosinophil granulocytes were measured with fluorescence flow cytometry. In the Sysmex WNR channel, basophil granulocytes were measured. Flow cytometry analysis was performed on the FACSCalibur™ system (Becton Dickinson, Heidelberg, Germany) using the BD Multitest™ IMK kit according to the manufacturer's instructions. BD Multitest™ IMK (CE/IVD) is a four-color direct immunofluorescence reagent for use in flow cytometry designed to determine in peripheral blood the major mature human lymphocyte sub-populations after lysis of erythrocytes, including the total number of T-lymphocytes (CD3+), B-lymphocytes (CD19+) and natural killer cells (CD3-CD56+ and/or CD56+) as well as helper/inducer (CD3+CD4+) and suppressor/cytotoxic (CD3+CD8+) T-lymphocyte subsets. Forward Scatter (FSC), Side Scatter (SSC), and fluorescence signals were determined for each cell. For lymphocyte analysis, two different panels were used. Sample tubes were equipped with the following antibody additions (Becton Dickinson): Tube 1:

CD3-FITC, CD8-PE, CD45-PerCP, CD4-APC; Tube 2: CD3-FITC, CD16-PE and CD56-PE, CD45-PerCP, CD19-APC. BD Multitest™ Lysing Solution was added to the tubes. After 15 minutes, incubation measurement was performed. CD45 was used for identification of total leucocyte population. Methods for differential blood cell count and for flow cytometric analysis for lymphocyte subsets as well as cortisol, IgA, IL-6, CRP, and 25-OH-vitamin D3 at the central laboratory are accredited according to DIN EN ISO 15189:2014. Analyses were performed in line with the guideline of the German Medical Association (RiliBÄK) according to stipulated internal and external quality controls.

For analyses of salivary oxytocin, cortisol, and IgA, we compared the area under the curve measuring the increase ($AUC_I$) in neuroendocrine levels as measurement sensitive for time differences between sample assessments (mean interval pre to post social interaction ± SD: 29.6 ± 4.7 minutes; post to 15 minutes post: 15.0 ± 2.5 minutes) [20].

**Psychophysiology data acquisition and analysis**

Electrodermal activity (EDA) and an electrocardiogram (ECG) were collected via acquisition module MP150 (Biopac Systems Inc., Goleta CA, USA). EDA was measured at a sampling rate of 1000 Hz from Ag/AgCl electrodes filled with isotonic electrolyte gel on the tenar and hypotenar of the left (non-dominant) hand. ECG data were measured at a sampling rate of 1000 Hz from Ag/AgCl electrodes filled with isotonic electrolyte gel and attached to the participants in a Lead-II formation with active electrodes positioned on the right clavicle and on the lower left rib and the ground electrode placed on the left clavicle. EDA and ECG data were analyzed with Acqknowledge 4.3 software (Biopac Systems Inc., Goleta CA, USA). Preprocessing of the EDA data included resampling of the waveform to a new waveform sample rate of 62.5 samples per second, smoothing (median value smoothing factor: 63), and applying a low-pass filter with a frequency cutoff of 1 Hz. EDA data were visually inspected for artifacts. Phasic components were derived from the tonic EDA before the skin conductance level (SCL) was assessed. The raw ECG data was first analyzed using an automated software detection algorithm and then visually inspected according to the software guidelines for detecting artifacts and abnormal beats.

**Detailed description of the calculated mixed analyses of variance (ANOVAs)**

The behavioral, psychophysiological, and functional connectivity data were analyzed by calculating 2 x 2 mixed-measure ANOVAs, i.e. repeated measure ANOVAs that further included the between-subject factor group (HL vs. LL).[21] While each participant was assigned to either the HL or the LL group, all participants underwent all factor levels of the within-subject factors. For all analyses, main effects of group and main effects of the respective within-subject factors were calculated as well as the interactions between group and the within-subject factor (i.e. whether different effects of the within-subject factors were observed depending on the group membership of the participants). In detail, to analyze the behavioral and the functional connectivity data of the trust game, we tested for main effects of group and the sociality condition (trust game vs. risk game) and for the interaction of group and sociality condition. For self-reported affect and mood and the interpersonal distances, main effects of group and time (before vs. after the social interaction) were calculated in addition to the interaction of group with time. Likewise, main effects of group and time (baseline vs. during social interaction) and the interaction of group and time were analyzed for the SCL and heart rate (beats per minute, BPM).

### Correlation, mediation and moderation analyses

We correlated parameter estimates of trust-specific brain activity and connectivity (trust game > risk game) with those variables that were associated with loneliness (main effects of group or interactions with group), i.e. the interpersonal trust as trait variable, the mean investment across trust and risk game trials, the interpersonal distance across time in the ideal and uncomfortable condition, the positive affect (increase from pre to post completion of the positive social interaction task as well as separately for pre and post ratings), and the oxytocinergic responsiveness as calculated by the $AUC_I$. To explore the relationship of interpersonal trust with behavioral data, we also correlated the self-reported interpersonal trust with the above-mentioned variables.

To examine whether group effects (see the above mentioned variables as well as reduced neural discrimination between trust and risk game, reduced trust-related connectivity, and immunology) might be driven by or interact with psychiatric symptomatology, mediation and moderation analyses were carried out using the PROCESS macro v3.4 for SPSS.[22] BDI, LSAS, and CTQ scores were used as mediator and moderator variables and group as independent variable. Further moderation analyses were conducted to examine potential effects of sex. Moderator variables were mean centered. For mediation analyses, 10,000 bootstrap samples were used.

### Missing data

Blood and saliva samples were not available for all participants due to problems in sample assessment (missing values for full blood count: $n = 14$, oxytocin, CRP, IL-6, and vitamin D blood: $n = 3$, salivary oxytocin: $n = 2$, salivary cortisol: $n = 11$, salivary IgA: $n = 34$). Samples with concentrations less than the minimum detection limit were included into the analysis using the detection limit as data values. For missing individual time intervals between saliva samples after completion of the social interaction task ($n$ = 16), time intervals of fifteen minutes (i.e. the planned time interval) were used for the computation of the area under the curve. EDA and ECG data were missing for four (EDA) and six (ECG) participants, respectively. Only a subsample of 50 participants completed the Pittsburgh Sleep Quality Index.[7]

**Supplementary Analyses**

**Associations of loneliness with immunology**

Notably, while the immunological responsiveness to the social interaction (i.e. increase in IgA concentrations) did not significantly differ between groups ($t_{(44)} = 0.51$, $P = 0.610$, 95 % CI of mean $AUC_I$ difference between groups: -0.34 to 0.57; see **Figure S1A**), HL participants showed lower absolute numbers and percentage of lymphocytes (all *t*s < -2.42, all *P*s < 0.019, 95 % CI of group difference in absolute numbers of lymphocytes: -0.63 to -0.06 G/l; percentage of lymphocytes: -8.15 to -0.78 %) due to lowered total CD3 T-lymphocytes ($t_{(63)} = -2.52$, $P = 0.014$, $d = -0.63$, 95 % CI of group difference: -532.00 to -61.51 per µl) and regulatory CD3 and CD8+ T-lymphocytes ($t_{(63)} = -3.15$, $P = 0.003$, $d = -0.78$, 95 % CI of group difference: -250.18 to -55.86 per µl; **see Figure S1B**) in the baseline full blood count, indicating a weakened immune system in HL participants.

**Further associations of loneliness with mood**

The positive social interaction task caused an increase in happiness as assessed via a VAS [main effect of time: $F_{(1,77)} = 6.45$, $P = 0.013$, $\eta_p^2 = 0.08$, 95 % confidence interval (CI) of increase: 0.82 to 6.60], while a decrease in negative affect was evident for rated sadness (main effect of time: $F_{(1,77)} = 7.81$, $P = 0.007$, $\eta_p^2 = 0.09$, 95 % CI of decrease: -6.16 to -0.95) and the following scales of the POMS: depression ($F_{(1,77)} = 7.99$, $P = 0.006$, $\eta_p^2 = 0.09$, 95 % CI of decrease: -1.66 to -0.27), fatigue ($F_{(1,77)} = 67.53$, $P < 0.0001$, $\eta_p^2 = 0.47$, 05 % CI of decrease: -6.67 to -4.08), and anger ($F_{(1,77)} = 11.25$, $P = 0.001$, $\eta_p^2 = 0.13$, 95 % CI of decrease: -0.79 to -0.20). Ratings on the VAS regarding anger, anxiety, state loneliness, overall arousal, and negative affect as measured with the

PANAS were unaffected by the positive social interaction task (all *P*s > 0.069). HL participants, however, differed from LL participants in almost all measurements of mood pooled over time except for the positive affect as measured with the PANAS, fatigue as measured with the POMS, and the overall arousal. Specifically, HL participants reported reduced positive mood (main effect of group for VAS happiness and POMS vigor; all *P*s < 0.040, 95 % CI of group difference for VAS happiness: -21.23 to -6.94; POMS vigor: -7.36 to -0.19), whereas negative affect (anger, anxiety, sadness, state loneliness, PANAS negative, POMS depression, and POMS anger; all *P*s < 0.047; 95 % CI of group difference for state anger: 0.38 to 5.99; anxiety: 0.56 to 9.29; sadness: 6.68 to 17.92; state loneliness: 11.10 to 24.18; PANAS negative: 0.37 to 2.58; POMS depression: 1.16 to 6.64; POMS anger: 0.00 to 2.00) and BPM ($F_{(1,70)} = 5.35$, $P = 0.024$, $\eta_p^2 = 0.07$, 95 % CI of group difference: 1.16 to 8.91 BPM) were increased. An interaction effect of time (before versus after completion of the positive social interaction task) with group (HL versus LL) indicated a greater reduction of sadness in HL participants ($F_{(1,77)} = 4.07$, $P = 0.047$, $\eta_p^2 = 0.05$), but post-hoc *t*-tests showed still significantly greater sadness ratings of HL participants compared to controls after completion of the task [pre: $t_{(41.02)} = 4.39$, after Bonferroni-correction ($P_{cor}$) $P_{cor} = 0.0001$, $d = 0.99$, 95 % CI of group difference: 8.03 to 21.74; post: $t_{(39.30)} = 3.59$, $P_{cor} = 0.002$, $d = 0.81$, 95 % CI of group difference: 4.24 to 15.18]. Notably, the interaction effect appears to be driven by a bottom effect in the LL participants, as self-rated sadness before the interaction was minimal (mean sadness ratings before the interaction in LL participants ± SD: 1.60 ± 4.20; HL participants: 16.49 ± 20.79; mean sadness ratings after the interaction in LL participants ± SD: 0.60 ± 2.22; HL participants: 10.31 ± 16.75). Results revealed no further group differences for self-reported measurements or ratings by the participants, ratings by the experimenter, SCL, or measurements of baseline cortisol levels and increases as measured using the $AUC_I$ (all *P*s > 0.054).

### Further analyses of the fMRI trust game

Whole-brain analyses confirmed the involvement of the ROIs in the trust game across groups. The amygdala [left: -18, -6, -16, $t_{(58)} = 8.95$, $P < 0.001$ after family-wise error correction (FWE) for the number of voxels across the whole brain; right: 22, -6, -12, $t_{(58)} = 11.35$, FWE-corrected $P < 0.001$], AI (left: -30, 14, -16, $t_{(58)} = 7.91$, FWE-corrected $P < 0.001$; right: 26, 18, -16, $t_{(58)} = 7.57$, FWE-corrected $P < 0.001$), mPFC (left: -2, 46, -12, $t_{(58)} = 10.44$, FWE-corrected $P < 0.001$; right: 2, 46, -12, $t_{(58)} = 10.79$, FWE-corrected $P < 0.001$ and 2, 52, 22, $t_{(58)} = 10.04$, FWE-corrected $P < 0.001$), and TPJ (left: -56, -60, 28, $t_{(58)} = 7.37$, FWE-corrected $P < 0.001$; right: 58, -54, 24, $t_{(58)} = 8.17$, FWE-corrected $P < 0.001$) showed increased activity during the trust game compared to the risk game. No significant effects across groups were observed for the NAcc.

An exploratory whole-brain analysis revealed a frontoparietal network that was active during trust evaluations compared to age ratings (trust game > age human; see **Table S3**). Interestingly, several trust-associated brain regions including the AI showed no significant activation when age ratings were included in a differential contrast (i.e. [(trust game > age human) > (risk game > age computer)]), suggesting that the social content is a necessary component for interpersonal trust. In fact, no significant differences between LL and HL participants were observed when comparing brain activity during the trust game with the age ratings [i.e. trust game > age human and (trust game > age human) > (risk game > age computer); all FWE-corrected $P$s $\geq 0.129$].

While no significant group differences were observed when calculating exploratory whole-brain analyses, Bayesian inference analyses provide strong evidence for increased trust-associated brain activity (trust game > risk game) in lonely participants in the right cuneus [4, -74, 34, log odds

Bayes factor for greater activity in HL participants versus no group differences or decreased activity in HL participants (logBF) = 3.51], the right lobule VI of cerebellar hemisphere (14, -74, -14, logBF = 3.26), the precuneus (0, -62, 50, logBF = 3.06), and the right superior temporal gyrus (46, -10, -4, logBF = 3.05; see **Figure S2**).

### Further analyses of connectivity

We calculated additional post-hoc analyses to enable a better understanding of the observed group differences in functional connectivity of the left AI with the occipitoparietal cluster. As the observed reduced connectivity in HL participants during the trust game compared to LL participants might not automatically indicate a weaker positive connectivity but could also be based on a stronger negative connectivity (i.e. stronger inhibitory effects) in HL participants, we tested whether trust game connectivity values of each group significantly differed from zero. Results revealed a significant positive connectivity in the LL group ($t_{(27)} = 3.53$, $P = 0.002$, $d = 0.67$, 95 % CI of connectivity effect size: 0.08 to 0.30) whereas connectivity in the HL sample was not significantly different from zero ($t_{(30)} = -1.25$, $P = 0.222$, 95 % CI of connectivity effect size: -0.22 to 0.05), indicating that the reported reduced connectivity was not based on a stronger negative connectivity in HL individuals.

Further exploratory seed-to-voxel analyses were calculated with those ROIs serving as seed regions whose task-dependent activity was not significantly associated with loneliness. Analyses revealed marginally significant reduced trust-associated connectivity of the left TPJ with a striatal cluster in HL compared to LL participants (-14, -06, 14, $k = 83$, $t_{(57)} = 5.42$, FWE-corrected $P = 0.050$ on cluster level). Correspondingly, analyses revealed reduced trust-associated connectivity of the left

NAcc with a temporoparietal cluster in HL participants (58, -56, 12, $k$ = 89, $t_{(57)}$ = 4.67, FWE-corrected $P$ = 0.035 on cluster level). As reported for the blunted AI connectivity (see main text), functional connectivity during the trust game was significantly diminished in HL participants compared to the LL controls (left TPJ: $t_{(57)}$ = -2.79, $P_{cor}$ = 0.028, $d$ = -0.73, 95 % CI of group difference: -0.40 to -0.07; left NAcc: $t_{(57)}$ = -2.99, $P_{cor}$ = 0.016, $d$ = -0.78, 95 % CI: -0.19 to -0.04), while connectivity during the risk game did not differ between groups (all $t$s < 1.62, all Bonferroni-corrected $P$s > 0.447). Likewise, LL participants showed significantly increased functional connectivity during the trust game compared to the risk game (left TPJ: $t_{(27)}$ = 4.30, $P_{cor}$ < 0.001, $d$ = 0.68, 95 % CI of increase: 0.11 to 0.31; left NAcc: $t_{(27)}$ = 3.91, $P_{cor}$ = 0.002, $d$ = 0.74, 95 % CI: 0.05 to 0.17), whereas functional connectivity decreased in HL participants (left TPJ: $t_{(30)}$ = -3.32, $P_{cor}$ = 0.009, $d$ = -0.47 3, 95 % CI of decrease: -0.24 to -0.06; left NAcc: $t_{(30)}$ = -2.59, $P_{cor}$ = 0.059, $d$ = -0.45, 95 % CI: -0.12 to -0.01). No further significant differences in trust-associated connectivity were observed between groups.

**Further moderation analyses of group effects**

Depressive symptoms moderated the group effects on the mean invested money across trust and risk game trials (significant interaction term of group and BDI scores; $B$ = 0.48, $t_{(61)}$ = 2.45, $P$ = 0.017, 95 % CI: 0.09 to 0.86). The Johnson-Neyman technique identified a complex interaction of depressive symptoms and loneliness such that less money was invested by HL individuals if the BDI scores were 2.80 or lower ($B$ = -1.17, $t_{(61)}$ = -2.00, $P$ = 0.050, 95 % CI: -2.35 to 0.00), while more money was invested if the BDI scores were higher than 19.42 ($B$ = 6.74, $t_{(61)}$ = 2.00, $P$ = 0.050, 95 % CI: 0.00 to 13.47). Furthermore, depressive symptoms moderated the effect of loneliness on the immune system [significant interaction terms of group (HL versus LL) and BDI scores for absolute numbers of lymphocytes; $B$ = -0.12, $t_{(64)}$ = -2.81, $P$ = 0.007, 95 % CI: -0.21 to -0.03, absolute CD3 T-lymphocytes; $B$ = -126.59, $t_{(61)}$ = -3.36, $P$ = 0.001, 95 % CI: -201.93 to -51.25, and total regulatory CD3 and CD8+ T-lymphocytes; $B$ = -55.53, $t_{(61)}$ = -3.64, $P$ = 0.001, 95 % CI: -85.99 to -25.06]. For all measurements, the Johnson-Neyman technique showed that HL individuals with more depressive symptoms exhibited a decreased level of lymphocytes (BDI scores of 2.38 or higher for total lymphocytes: $B$ = -0.30, $t_{(61)}$ = -2.00, $P$ = 0.050, 95 % CI: -0.59 to 0.00; BDI scores of 1.99 or higher for total CD3 T-lymphocytes: $B$ = -241.41, $t_{(61)}$ = -2.00, $P$ = 0.050, 95 % CI: -482.82 to 0.00; BDI scores of 1.07 or higher for CD3 and CD8+ T-lymphocytes: $B$ = -104.43, $t_{(61)}$ = -2.00, $P$ = 0.050, 95 % CI: -208.86 to 0.00), whereas the association of loneliness with immune parameters was not significant when no depressive symptoms were present, indicating that depressive symptomatology might enhance the negative association of loneliness and immune system parameters. Likewise, the effects of loneliness on the absolute number of lymphocytes was more pronounced with stronger social anxiety symptoms measured with the LSAS ($B$ = -0.02, $t_{64)}$ = -2.01, $P$ = 0.049, 95 % CI: -0.05 to -0.0001). The Johnson-Neyman technique showed that the association between loneliness and the absolute number of lymphocytes was significant for individuals with social anxiety scores of 8.62 or higher ($B$ = -0.30, $t_{(64)}$ = -2.00, $P$ = 0.050, 95 % CI: -0.61 to 0.00).

Notably, childhood maltreatment as measured by the CTQ significantly moderated the group effect on trust-related NAcc activity ($B$ = 0.01, $t_{(55)}$ = 2.11, $P$ = 0.039, 95 % CI: 0.001 to 0.02) and on the affective responsiveness to the positive social interaction paradigm as measured by the PANAS positive ($B$ = 0.18, $t_{(75)}$ = 2.19, $P$ = 0.032, 95 % CI: 0.02 to 0.35). In both cases, the negative effect of loneliness was enhanced when CTQ scores were low (significant regions as identified by the

Johnson-Neyman technique for NAcc activity: CTQ scores of 36.98 or lower, $B = -0.11$, $t_{(55)} = -2.00$, $P = 0.050$, 95 % CI: -0.21 to 0.00; for PANAS positive: scores of 39.82 or lower, $B = -2.07$, $t_{(75)} = -1.99$, $P = 0.050$, 95 % CI: -4.15 to 0.00), indicating that the impact of loneliness on the rewarding effects of social interactions became less important for participants suffering from more severe childhood trauma. We observed no further significant moderations for any of the reported main results (all *P*s of interactions between group and moderators > 0.063).

**Analysis of the trust game decisions while playing the role of the trustee**

We analyzed participants' decisions to keep all the money or to share it with the investor while playing the trust game in the role of the trustee during the screening session. Across groups, participants decided to share in 6.97 (SD: 3.05) out of 10 trials. HL participants differed neither in the total number of decisions to share across all possible investments ($t_{(63)} = -0.16$, $P = 0.875$, 95 % CI of group difference: -1.65 to 1.41) nor in the proportion of share decisions calculated separately for each possible investment as analyzed using chi-square tests (all *P*s > 0.260) except for the decisions following an investment of 3 Euro ($\chi^2_{(1)} = 4.48$, $P = 0.034$; 79.41 % of HL participants versus 54.84 % of LL participants decided to share).

### Supplementary Figures

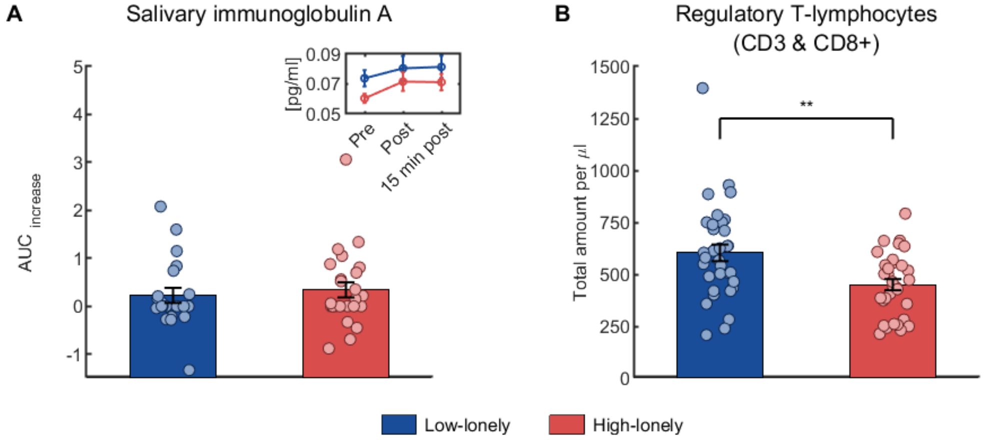


**Figure S1. Immune system reactivity to the positive social interaction paradigm and baseline regulatory T-lymphocytes count.** Across groups, the area under the curve measuring the increase ($AUC_{increase}$) in salivary immunoglobulin A (IgA) concentrations in response to the positive social interaction was significantly larger than zero (**A**). The inlay displays the mean salivary IgA concentrations for each time point. While high-lonely participants did not differ in their immune reactivity as measured with salivary IgA concentrations, loneliness was associated with altered baseline blood parameters. High-lonely participants showed a significantly reduced amount of regulatory CD3 and CD8+ T-lymphocytes (**B**), indicating a weakened immune system in high-lonely participants. All bars represent group means. Error bars indicate standard errors of the mean. Dots are jittered for purposes of presentation. ** $P < 0.01$.

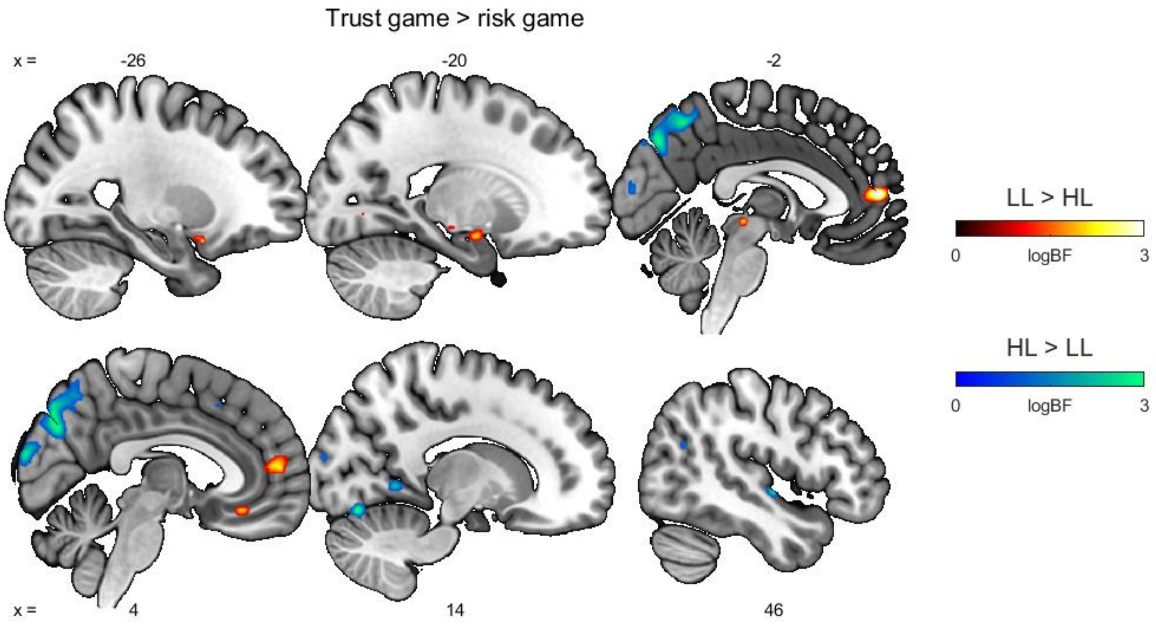


**Figure S2. Results of the Bayesian inference analyses for group differences in trust-associated brain activity.** Our data provide strong evidence with log odds Bayes factors (logBF) ≥ 3 for reduced trust-associated brain activity in lonely participants (LL > HL) in the left anterior insula (-26, 10, -18, logBF = 3.28) and the medial prefrontal cortex (0, 52, 10, logBF = 3.62), whereas strong evidence for increased trust-associated brain activity (HL > LL) was found for the right cuneus (4, -74, 34, logBF = 3.51), the right lobule VI of cerebellar hemisphere (14, -74, -14, logBF = 3.26), the precuneus (0, -62, 50, logBF = 3.06), and the right superior temporal gyrus (46, -10, -4, logBF = 3.05). Moderate evidence (i.e. positive evidence according to[23]) for group differences was found for various further clusters including the left amygdala (LL > HL: -20, -6, -14, logBF = 2.16). Only voxels with moderate evidence for at least small group differences are presented (logBF ≥ 1, *d* ≥ 0.2). Abbreviations: HL, high-lonely; LL, low-lonely; logBF, log odds Bayes factor.

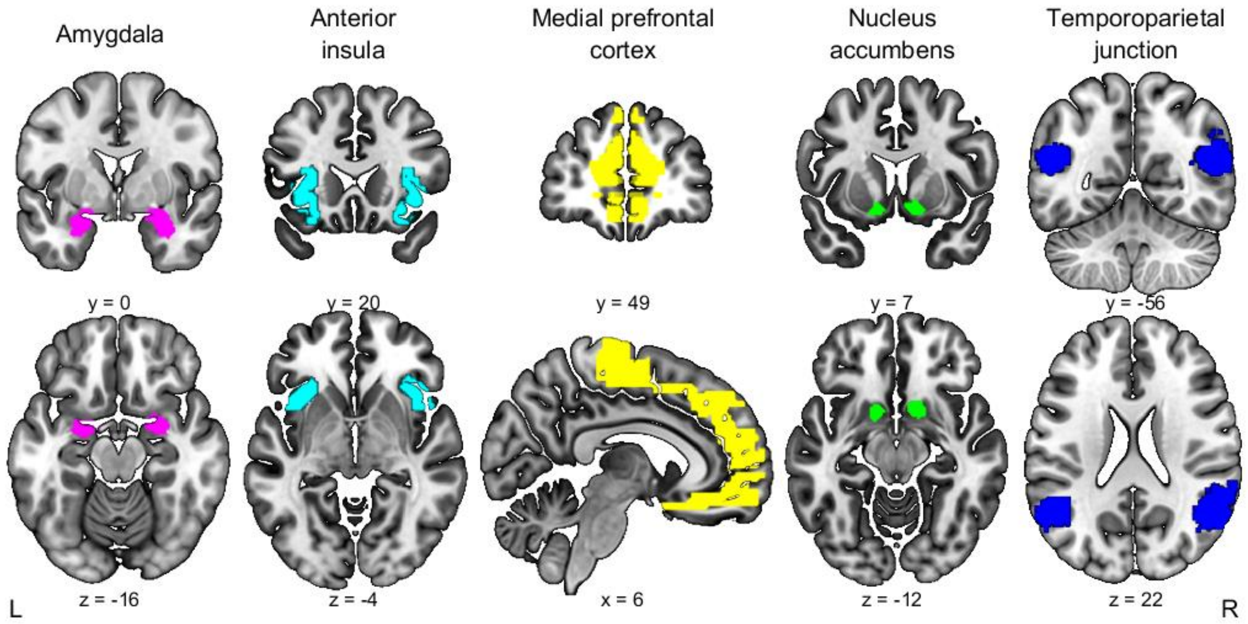


**Figure S3. Masks of the regions of interest.** The regions of interest that were used for the small volume correction were independently defined by using anatomical masks of the amygdala, anterior insula, medial prefrontal cortex, and the nucleus accumbens as implemented in the Wake Forest University PickAtlas.[15,16] The temporoparietal junction was independently created using association test maps from neurosynth (https://neurosynth.org/; search term: temporoparietal junction). Abbreviations: L, left; R, right.

**Supplementary Tables**

**Table S1.** Sociodemographic variables

| **Sociodemographic measurements** | **High-lonely** ($n$ = 42) | **Low-lonely** ($n$ = 40) | ***P*** |
|---|---|---|---|
| **Age (years), *M* (*SD*)** | 26.5 (6.8) | 27.1 (8.2) | 0.73 |
| **Education (years), *M* (*SD*)** | 14.9 (5.3) | 15.8 (3.9) | 0.39 |
| **Monthly salary, *n* (%)** | | | 0.44 |
| 0 to 500 € | 12 (28.6) | 12 (30.0) | |
| 501 to 1,000 € | 11 (26.2) | 16 (40.0) | |
| 1,001 to 1,500 € | 10 (23.8) | 6 (15.0) | |
| 1,501 to 2,000 € | 5 (11.9) | 1 (2.5) | |
| 2,001 to 2,500 € | 2 (4.8) | 1 (2.5) | |
| 2,501 to 3,000 € | 1 (2.4) | 1 (2.5) | |
| More than 3,000 € | 1 (2.4) | 3 (7.5) | |
| **BMI, *M* (*SD*)** | 25.1 (6.5) | 24.1 (3.5) | 0.41 |
| **Smoker, *n* (%)** | 7 (16.7) | 3 (7.5) | 0.21 |

*Notes.* Abbreviations: BMI, body mass index.

**Table S2.** Full blood count

| **Blood parameter** | **High-lonely** | **Low-lonely** | ***T*** | ***P*** |
|---|---|---|---|---|
| C-reactive protein [mg/l] | 2.7 (9.2) | 2.2 (3.0) | 0.33 | 0.75 |
| Interleukin-6 [pg/ml] | 2.4 (2.0) | 1.8 (0.9) | 1.60 | 0.12 |
| 25-hydroxyvitamin D [ng/ml] | 24.3 (12.8) | 27.6 (8.0) | -1.39 | 0.17 |
| Leukocytes [G/l] | 7.0 (2.2) | 6.8 (1.4) | 0.30 | 0.77 |
| Erythrocytes [T/l] | 5.0 (0.4) | 4.9 (0.4) | 1.65 | 0.10 |
| Oxytocin [pg/ml] | 3.7 (0.9) | 3.7 (1.1) | 0.13 | 0.90 |
| Thrombocytes (fl) [G/l] | 256.4 (54.2) | 268.6 (51.1) | -0.95 | 0.35 |
| Erythroblasts [/100 leukocytes] | 0.003 (0.02) | 0.024 (0.06) | -1.91 | 0.06 |
| Erythroblasts absolute [G/l] | 0.0003 (0.002) | 0.0015 (0.004) | -1.72 | 0.09 |
| Neutrophil granulocytes absolute [G/l] | 4.3 (1.9) | 3.9 (1.1) | 1.24 | 0.22 |
| **Lymphocytes absolute [G/l]** | **1.9 (0.5)** | **2.2 (0.6)** | **-2.45** | **0.02** |
| Monocytes absolute [G/l] | 0.6 (0.2) | 0.5 (0.1) | 0.98 | 0.33 |
| Eosinophil granulocytes absolute [G/l] | 0.2 (0.1) | 0.2 (0.1) | -0.95 | 0.35 |
| Basophil granulocytes absolute [G/l] | 0.05 (0.02) | 0.05 (0.02) | -0.53 | 0.60 |
| Neutrophil granulocytes [%] | 60.3 (8.9) | 56.1 (8.4) | 1.99 | 0.05 |
| **Lymphocytes [%]** | **28.5 (7.7)** | **33.0 (7.5)** | **-2.42** | **0.02** |
| Monocytes [%] | 8.1 (1.7) | 7.7 (2.0) | 1.04 | 0.30 |
| Eosinphil granulocytes [%] | 2.4 (1.8) | 2.5 (1.6) | -0.41 | 0.69 |
| Basophil granulocytes [%] | 0.7 (0.4) | 0.7 (0.3) | -0.25 | 0.81 |

| | | | | |
|---|---|---|---|---|
| B-cells CD19+ [%][a)] | 13.0 (4.8) | 11.3 (3.5) | 1.58 | 0.12 |
| B-cells CD19+ absolute [/µl][a)] | 248.3 (123.5) | 249.0 (88.9) | -0.03 | 0.98 |
| NK-cells CD16 & CD56+ [%][a)] | 14.6 (6.4) | 13.9 (5.4) | 0.47 | 0.64 |
| NK-cells CD16 & CD56+ absolute [/µl][a)] | 272.1 (124.9) | 305.3 (147.9) | -0.98 | 0.33 |
| T-lymphocytes total (CD3+) [%][a)] | 71.2 (6.5) | 73.3 (6.0) | -1.40 | 0.17 |
| **T-lymphocytes total (CD3+) absolute [/µl]**[a)] | **1,359.0 (408.5)** | **1,655.8 (530.7)** | **-2.52** | **0.01** |
| T-helper cell CD3 & CD4+ [%][a)] | 44.0 (7.0) | 42.6 (6.4) | 0.80 | 0.43 |
| T-helper cell CD3 & CD4+ absolute [/µl] [a)] | 848.8 (310.7) | 966.3 (362.8) | -1.40 | 0.17 |
| Regulatory T-cell CD3 & CD8+ [%][a)] | 24.1 (6.2) | 26.7 (5.7) | -1.76 | 0.08 |
| **Regulatory T-cell CD3 & CD8+ absolute [/µl]**[a)] | **450.3 (152.0)** | **603.4 (230.7)** | **-3.15** | **0.003** |
| CD4/CD8 ratio[a)] | 2.0 (0.9) | 1.8 (0.5) | 1.92 | 0.06 |

*Notes.* Values are mean and SD. Bold markings indicate significant group differences. [a)]*N* = 65.

Abbreviations: CD, cluster of differentiation; NK-cells, natural killer cells.

**Table S3.** Whole-brain findings across groups (thresholded at $P < 0.001$)

| Region | Right/left | Cluster size (voxel) | Peak *T* | MNI coordinates | | |
|---|---|---|---|---|---|---|
| | | | | x | y | z |
| **Trust game > risk game** | | | | | | |
| Midcingulate gyri | bil. | 2,218 | 13.56 | 2 | -54 | 32 |
| Hippocampus | R | 2,758 | 11.89 | 20 | -6 | -14 |
| Superior medial orbitofrontal gyri | bil. | 6,185 | 10.85 | 0 | 44 | -14 |
| Angular gyrus | R | 1,451 | 8.17 | 58 | -54 | 24 |
| Angular gyrus | L | 665 | 8.04 | -54 | -60 | 30 |
| Insula | R | 436 | 7.57 | 26 | 18 | -16 |
| Fusiform gyrus | R | 136 | 7.11 | 42 | -48 | -18 |
| Middle temporal gyrus | L | 518 | 7.00 | -56 | -24 | -12 |
| Inferior frontal gyrus (opercularis) | R | 532 | 5.98 | 42 | 14 | 28 |
| Inferior temporal gyrus | L | 52 | 5.43 | -40 | -48 | -18 |
| **Trust game > age human** | | | | | | |
| Midcingulate gyri | bil. | 8,332 | 12.12 | -2 | -22 | 38 |
| Superior frontal gyri | bil. | 12,031 | 11.66 | -12 | 30 | 56 |
| Inferior parietal gyrus | L | 4,597 | 11.65 | -52 | -58 | 44 |
| Inferior orbitofrontal gyrus | L | 662 | 7.36 | -34 | 16 | -18 |
| Middle frontal gyrus | R | 393 | 5.87 | 38 | 24 | 42 |

| | | | | | | |
|---|---|---|---|---|---|---|
| **Age human > age computer** | | | | | | |
| Precuneus | bil. | 1,702 | 12.05 | 2 | -54 | 36 |
| Angular gyrus | R | 5,718 | 10.69 | 54 | -52 | 24 |
| Superior medial frontal gyri | bil. | 4,050 | 10.10 | 2 | 56 | 14 |
| Hippocampus | L | 319 | 9.10 | -18 | -10 | -14 |
| Angular gyrus | L | 528 | 8.56 | -56 | -58 | 32 |
| Middle temporal gyrus | L | 200 | 6.05 | -58 | -12 | -14 |
| Midcingulate gyri | bil. | 151 | 5.79 | 2 | -20 | 38 |
| Inferior frontal gyrus (triangularis) | R | 118 | 5.43 | 54 | 32 | 6 |
| **(trust game > age human) > (risk game > age computer)** | | | | | | |
| Inferior temporal gyrus | L | 7,791 | 10.23 | -46 | -64 | -8 |
| Posterior cingulate gyri | bil. | 1,280 | 9.65 | 0 | -34 | 32 |
| Superior motor area | bil. | 14,007 | 9.54 | -2 | 20 | 50 |
| Angular gyrus | R | 7,177 | 9.41 | 32 | -54 | 42 |
| Nucleus caudatus | R | 2,818 | 8.81 | 12 | 6 | 10 |
| Anterior cingulate gyri | bil. | 159 | 6.04 | -2 | 6 | 28 |
| Gyrus rectus | bil. | 105 | 5.73 | 0 | 18 | -16 |

*Notes.* Only cluster with FWE-corrected *P*s < 0.05 on peak level are listed. Abbreviations: bil., bilateral; L, left; MNI, Montreal Neurological Institute; R, right.

**Table S4.** Psychiatric symptomatology and social network description

| **Psychiatric measurements** | **High-lonely** (*n*=42) | **Low-lonely** (*n*=40) | ***T*** | ***P*** |
|---|---|---|---|---|
| **UCLA-L**[a)] | **57.0 (5.4)** | **23.8 (1.3)** | **38.76** | **< 0.0001** |
| **BDI**[b)] | **6.6 (6.8)** | **2.0 (2.3)** | **4.15** | **< 0.001** |
| **LSAS**[c)] | **18.6 (15.9)** | **9.3 (9.6)** | **3.25** | **0.002** |
| **CTQ**[d)] | **38.9 (10.3)** | **31.9 (15.8)** | **2.38** | **0.020** |
| **PSQI**[\|e)] | **8.4 (2.6)** | **6.3 (1.7)** | **3.19** | **0.002** |
| **Social Network Size**[f)] | | | | |
| Number of social roles | **5.0 (1.5)** | **6.1 (1.4)** | **3.36** | **0.001** |
| Total number of persons within the network | **15.4 (7.1)** | **21.9 (7.5)** | **4.07** | **< 0.001** |

*Notes.* Values are mean and SD. [a)]Participants were pre-stratified and assigned to the high- or low-lonely group using the UCLA Loneliness Scale (UCLA-L). High-lonely participants had a score equal or above 50 while low-lonely participants had a score equal or below 25; [b)]Depressive symptoms were measured with the Beck Depression Inventory, Version II (BDI); [c)]Social anxiety was assessed with the Liebowitz Social Anxiety Scale (LSAS); [d)]Childhood traumata were measured using the Childhood Trauma Questionnaire (CTQ); [e)]Fifty participants (high-lonely: *n* = 29) completed the Pittsburgh Sleep Quality Index to assess sleep quality; [f)]Social network was characterized using the Social Network Index assessing the number of diverse social roles (maximum twelve roles) and the total number of people to whom the participants talk to regularly within all of their roles.
